# THE BILE ACID RECEPTOR TGR5 MODULATES HIPPOCAMPAL NEURAL STEM CELL STATE AND AGEING

**DOI:** 10.64898/2026.10.08.757807

**Authors:** Devanshu Kaushik, Prachi Dugam, Gourav Sarkar, Boda Arun Kumar, Sandip Patel, Sai Prasad Pydi, Prem N. Yadav

## Abstract

Ageing locks hippocampal neural stem cells (NSCs) into deeper quiescence and inflammation-driven senescence, which are also associated with impaired cognition. The bile-acid receptor TGR5, a member class I G-protein coupled receptors (GPCRs), is expressed in cultured hippocampal NSCs and in SOX2-positive cells of the adult sub granular zone, yet it’s role in NSC homeostasis is unknown. In this study we show that TGR5 selective agonist, TC-G 1005, expanded the NSC pool without forcing neuronal differentiation. Primary NSCs treatment with TC-G 1005 increased BrdU incorporation and PCNA in a concentration-dependent manner, and both responses were lost after TGR5 knockdown. TGR5 agonism in differentiating cultures increased nestin, left doublecortin (DCX) unchanged and suppressed MAP2 expression, preserving stemness rather than neuronal commitment. The agonist also released BMP4-imposed quiescence, lowering FOXO3a and restoring phospho-EGFR and ASCL1, and reopened autophagic flux, with less p62 and LC3-II, fewer LAMP2 puncta and more Beclin-1. Furthermore, activation of TGR5 suppressed IL-6 induced p53 and p16 expression, restored proliferation and contracted the inflammatory transcriptome by more than 90%, including Cdkn2a, Cdkn2b and Cdk6. In 18-month-old mice, brain-penetrant TC-G 1005 (5 mg/kg, subcutaneously, twice daily for 28 days) expanded SOX2-positive and proliferating NSCs, reduced p16, recovered ASCL1 and improved spatial memory, without changing locomotion or generating DCX-positive neurons. Dentate gyrus-restricted TGR5 knockdown abolished these cellular effects and the rescue of spatial and social discrimination. Hippocampal proteomic analysis revealed that TC-G 1005 treated aged mice occupied an intermediate, partially rejuvenated position between young and aged proteomes, with 86 proteins shared between ageing- and treatment-responsive sets, and mitochondrial and complement C1q modules shifted toward youthful levels. TGR5 activation therefore relieves both quiescence and senescence in the aged neurogenic niche without driving neuronal differentiation. Collectively, these results suggest that TGR5 could be targeted to preserve cognition in ageing by rejuvenating the hippocampal stem-cell niche.

## INTRODUCTION

Ageing is accompanied by progressive deficits in learning, memory and cognitive flexibility that track hippocampal dysfunction (1). The dentate gyrus sustains a neurogenic niche in the sub granular zone, where radial glia-like neural stem cells (NSCs) persist into adulthood, generate neurons and support spatial memory, mood regulation and stress resilience (2), (3), (4), (5). With age, the number of proliferative NSCs falls, fewer new neurons integrate, and the remaining stem cells become deeply quiescent (1), (6), (7). These changes coincide with impaired mitochondria, proteostatic stress and dysregulated growth-factor signalling (8), (9), (10). The unresolved question is which cell states produce this failure, and which of those states a drug can reverse.

Adult NSCs are maintained by quiescence, a reversible G_0_ arrest that protects the reservoir from exhaustion (11), (12). Quiescent cells are marked by increased FOXO3 and low ASCL1, with repressed EGFR signalling (13), (14), (15), (16), and this state can be imposed experimentally by BMP4 (17). Enhanced autophagolysosomal machinery content accumulation maintains NSCs quiescent, and restoring autophagic flux is associated with reactivation (16), (18), (17). Ageing deepens this quiescence state of NSCs and resist activation, yet cells that do activate can still behave like young stem cells, suggesting that the block is in state control rather than irreversible loss of competence (19). Notably, deeply quiescent NSCs in the mature brain express molecular hallmarks of stem-cell ageing (20). The pharmacological modulation of quiescence/activation state therefore appears a tangible strategy for niche repair. Senescence is commonly enforced by p53 and p16^INK4a^, and it is coupled to an inflammatory identity. Chronic low-grade inflammation, or inflammaging, contributes to cognitive decline. Interleukin-6 is sufficient to suppress hippocampal neurogenesis (21), and inflammatory signalling more broadly represses NSC self-renewal (22), (23). In the aged niche, inflammation may convert a reversible quiescent state into a senescence-like arrest from which proliferation is not readily restored. A useful intervention would therefore have to do two things: to release the lysosomal and FOXO3-linked quiescent programme, and interrupt p53–p16 senescence, without exhausting the pool by driving terminal neuronal commitment.

G protein-coupled receptors are highly druggable targets, and already known to tune NSC proliferation and differentiation through Wnt, Notch and Hedgehog signalling (24), (25), (26). Takeda G protein-coupled receptor 5 (TGR5, also GPBAR1) is a bile-acid receptor coupled to cAMP generation (27), (28). In the brain it is expressed in neurons and astrocytes, can act as a neurosteroid receptor, and its activation limits neuroinflammation (29), (30), (31). Outside the nervous system, bile-acid signalling through TGR5 activates intestinal stem cells and promotes epithelial regeneration (32). Whether TGR5 is present in hippocampal NSCs (_hi_NSC), and whether it controls their quiescence, autophagy, senescence or differentiation, is not known.

In this study we showed that TGR5 is expressed in cultured hippocampal NSCs and in SOX2-positive cells of the adult subgranular zone. Its selective agonist, TC-G 1005, increased proliferation but suppressed neuronal differentiation. TGR5 agonism also released BMP4-imposed quiescence, restored autophagy and suppressed IL-6-driven p53 and p16 expression. In 18-month-old mice, brain-penetrant TC-G 1005 expanded the SOX2-positive pool, reduced p16, recovered ASCL1 and improved spatial memory without generating DCX-positive neurons. These cellular and cognitive effects required hippocampal TGR5. Targeting this receptor may therefore rejuvenate the aged neurogenic niche by relieving quiescence and senescence, rather than by making new neurons.

## Material and methods

### Primary hippocampal neural stem cell culture

Isolation of mouse NSC/progenitors was done with modification as described previously (14). Neonatal mice pups (0-1 day old) were decapitated, and brains were rapidly removed into a 10-cm Petri dish containing 5 mL HBSS (Gibco 24020-117). The hippocampi were dissected under a stereomicroscope and transferred to 1.5 mL microcentrifuge tubes containing ∼500 µL cold HBSS. Tissue was centrifuged at 200 g for 1 min, the supernatant discarded, and the pellet incubated with 0.5% papain (400 µL) at 37 °C for 15–20 min with intermittent trituration. The digestion was stopped by adding neurobasal medium (Gibco; 12348-017) supplemented with B-27 plus supplement (Gibco; A35828) and N2 supplements (Gibco; 17502-048), followed by trituration (15 strokes) and centrifugation at 300 × g for 5 min at room temperature. Subsequently, the supernatant was discarded and the cell pellet was resuspended in Neurobasal medium and transferred to culture flasks supplemented with EGF (Genescript; Z-02691) and bFGF (Genescript; Z-03230) (20 ng/mL each). Growth factors were replenished daily until neurosphere formation was observed.

### Immunocytochemistry of hippocampal neural stem cells

NSC/Neurospheres were adhered onto poly-D-lysine–coated slides using a cytospin bucket and fixed with 2% paraformaldehyde (PFA) for 15 minutes. Cells were then permeabilized with 0.3% Triton X-100 in 1× PBS for 1 h at room temperature and blocked in 3% bovine serum albumin, 3% horse serum, and 0.3% Triton X-100 in PBS for 2 h. Samples were then incubated overnight at 4 °C with primary antibodies against Anti-MAP2, Anti-Nestin, Anti-BrdU, Anti-SOX2, Anti-Doublecortin, Anti-LAMP2, Anti-Ascl1, Anti-p16INK4a, Anti-TGR5, Anti-GFAP, and Anti-tGFP. After PBST washes, samples were incubated with Alexa Fluor 488- or 594- or 647-conjugated secondary antibodies (1:1000) and DAPI (2 µg/mL) for 1 h at room temperature. Slides were mounted using Vectashield antifade mounting medium, and fluorescence images were acquired using an Olympus BX61 FV1200-MPE microscope.

### Western Blotting

Western blotting was performed to assess protein expression as previously described (33). Frozen brain tissues and neurospheres were homogenised/lysed in lysis buffer comprising 25 mM Tris-HCl (pH 7.6), 150 mM NaCl, 0.5% NP-40, 1% sodium deoxycholate, and 0.1% SDS. Equal amounts of protein (25 μg) were separated on 8–12% SDS-PAGE and transferred onto a 0.2 μm PVDF membrane (Bio-Rad). Membranes were blocked with 5% bovine serum albumin in TBST for 2 h at room temperature and incubated overnight at 4 °C with primary antibodies against anti-p16 INK4a, anti-LC3β, anti-p53, Anti-beclin-1 (D40C5), Anti-PCNA (PC10), Anti-p16 INK4A (D7C1M), Anti-Ascl1 (E5S4Q), Anti-FoxO3a (75D8), Anti-SQSTM1/p62, Anti Total-EGFR, Anti-pEGFR (pTyr1068). After TBST washing, membranes were incubated with HRP-conjugated secondary antibodies (1:4000) for 2 h at room temperature. Signals were detected using enhanced chemiluminescence and imaged with an Amersham ImageQuant™ 800. Band intensities were quantified using ImageJ.

### Immunohistochemistry and Confocal Imaging

C57BL/6J mice were transcardially perfused, and brains were collected, fixed overnight in 4% paraformaldehyde, and cryoprotected in 30% sucrose for 48 h. Coronal brain sections (30 μm) were prepared using a cryostat (Thermo Scientific). Sections were washed in PBS, subjected to antigen retrieval in 10 mM sodium citrate buffer (pH 6.3) for 15 min and permeabilized with 0.5% Triton X-100 in PBS for 1 h at room temperature. After blocking with 3% bovine serum albumin, 3% horse serum, and 0.3% Triton X-100 in PBS for 2 h, sections were incubated overnight at 4 °C with primary antibodies against Anti-BrdU, Anti-SOX2, Anti-doublecortin, Anti-TGR5, Anti-Ascl1, Anti-p16INK4a, Anti RFP. Sections were then washed with PBST and incubated with Alexa Fluor 488- or 594- or 647-conjugated secondary antibodies (1:1000) and DAPI (2 µg/mL) for 1 hour at room temperature. After washing, sections were mounted on poly-D-lysine–coated slides using Vectashield antifade mounting medium. Fluorescent images were acquired using an Olympus BX61 FV1200-MPE microscope with a 40× and 60x oil objective with a 1 μm z-step. Total number of SOX2+ cells or SOX2+BrdU+ cell or SOX2+Ascl1+ cells were quantified by multiplying total number of cells to the total volume of the image

### Bulk-RNA Sequencing and data analysis

Vehicle, IL6 Treated and IL6+TC-G 1005 treated _hi_NSC were snap frozen in liquid nitrogen until processing. Sequencing was performed by Nucleome Informatic Private Limited, Hyderabad, India. Raw RNA-seq FASTQ files were processed using the nf-core/rnaseq v3.26.0 workflow, implemented in Nextflow v25.10.2 (34). Sequencing read quality was assessed using FastQC v0.12.1 before and after adapter and quality trimming. Read trimming was performed using Trim Galore! V2.1.0. The resulting reads were aligned to the Mus musculus GRCm39 reference genome using STAR v2.7.11b, with gene annotation based on the GENCODE mouse M38 GTF annotation. Transcript-level quantification was performed using Salmon v1.23.1. SAMtools v1.21 was used for alignment file processing, including sorting, indexing, and generation of alignment statistics. PCR and optical duplicate reads were identified using Picard MarkDuplicates v3.4.0, https://broadinstitute.github.io/picard/. The resulting raw, unnormalised gene-level count matrix was used for downstream differential expression analysis in R v4.3.3 using the DESeq2 package v1.40 (35). Differentially expressed genes (DEGs) were defined based on an absolute log2 fold-change threshold of 1.5 and a Benjamini–Hochberg-adjusted p-value threshold of 0.05. Gene sets used for pathway and gene-set enrichment analyses were obtained from the Mouse Molecular Signatures Database (MSigDB) (36) [3]. Data visualisation and downstream analyses were performed in R using appropriate accessory packages. Gene Ontology analysis of the DEGs was performed by the Gene Ontology (GO) Pathway and Molecular Function 2025 databases using enrichR (v3.4) package in R.

### Reverse Transcription (RT) and Quantitative PCR

Total RNA was extracted from primary neural stem cells using the NucleoSpin RNA XS kit (Macherey-Nagel, 740920.50) according to the manufacturer’s instructions. cDNA was analyzed by quantitative real-time PCR using TB Green chemistry (Takara RR086A) with primers specific for TGR5 and the housekeeping gene Actin. PCR was performed at 95 °C for 10 min, followed by 40 cycles of 95 °C for 10 s, 58 °C for 30 s, and 72 °C for 30 s. Actin (Forward: 5′-TGTTACCAACTGGGACGACA-3′; Reverse: 5′-CTGGGTCATCTTTTCACGGT-3′) was used for normalization, while TGR5 primers were Forward: 5′-TCAGTCTTGGCCTATGAGCGTCG-3′ and Reverse: 5′-TGGCTCTTCCTCGAAGCACTCG-3′. Relative gene expression was calculated using the 2^−ΔΔCt method.

### Hippocampal proteomics study

Hippocampus of young, old and old + TC-G 1005 treated mice was isolated and snap frozen in liquid nitrogen until further processing. Sample preparation and Mass Spectrometric Analysis of Peptide Mixtures were done at valerian chem private limited, Noida, Uttar Pradesh, India. Raw files were acquired and processed. Protein identification and quantification were performed using Proteome Discoverer version 3.2 (Thermo Fisher Scientific). Tandem mass spectra were searched against the UniProt Mouse (UP000000589) reference proteome database using the SEQUEST HT search engine. Oxidation of methionine and protein N-terminal acetylation were specified as variable modifications, whereas carbamidomethylation of cysteine was specified as a fixed modification. Peptide-spectrum matches were evaluated using a target–decoy strategy, and protein and peptide identifications were filtered at a false discovery rate (FDR) of ≤1%. Subsequent statistical analyses and data visualisation were conducted in R using RStudio (version 2016.08) and the relevant R packages.

### Animals

All in vivo experiments were conducted in accordance with the guidelines of the Institutional Animal Ethics Committee of the CSIR–Central Drug Research Institute, Lucknow, India. Young 2-4 months old (22-25 g) or and aged 15–18 months (25–28 g) male C57BL/6J mouse were used in this study. Animals were housed under a 12 h light/dark cycle at 22–25 °C and 50 ± 5% humidity, with ad libitum access to food and water.

### Drugs

TGR5 agonist TC-G 1005 (Medchem Express; HY-110173) was administered subcutaneously (s.c.) at a dose of 5 mg/kg for 28 days twice a day. Control animals were injected with vehicle solution containing 0.9% sodium chloride+0.1% DMSO. BrdU (Sigma; B5002) was injected intraperitoneally on last 3 days before sacrifice at 50 mg/kg body weight.

### Pharmacokinetic study

Pharmacokinetic profiling of TC-G 1005 was outsourced and conducted by Anthem Biosciences Private Limited, Bangalore, India. In brief, Adult healthy male C57BL/6 Mice (G1 to G5) were used for experimentation after a minimum 3 days of acclimation. Five sampling time points were selected: 0.5, 1, and 4, 8, 12 h post-dose. (n = 3 per time point). Non-fasted animals were administered with test item TC-G 1005 by subcutaneous route with a dose of 10 mg/kg body weight at a dose volume of 5 mL/kg body weight. Under mild isoflurane anesthesia, blood samples were collected into pre-labeled tubes containing anticoagulant (K2EDTA - 2 mg/mL blood) during the next 12 hours of post-dose. Collected blood samples were centrifuged at 6000 rpm, 4 °C for 10 minutes and plasma was separated and stored at -80 °C until analysis. After collection of blood samples, animals were euthanized with CO2 asphyxiation followed by exsanguination and brain was collected during the next 12 hours of post dose and other time points. Brain specimens were stored at -80 °C until analysis. LC-MS/MS (AB Sciex 3200) was used for the quantitative determination of TC-G 1005 in plasma and Brain tissues.

### AAV9 Viral preparation & Stereotaxy

TGR5 shRNA (5′-TCCTACCTCTACCTGGAAGTTT-3′) or scrambled shRNA (5′-CCTAAGGTTAAGTCGCCCTCG-3’) was cloned into the AAV9-EF1α-mCherry vector. AAV9 particles were packaged in HEK293 cells, purified using a discontinuous iodixanol gradient (Sigma-Aldrich), and concentrated to 1 × 10¹² genome copies/mL. A total of 0.5 μL viral suspension was bilaterally injected into the dentate gyrus (AP: -2.1 mm, ML: 1.3 mm, DV: 1.9 mm) using a motorized stereotaxic system (Stoelting, USA). Mice were anesthetized with ketamine/xylazine (90/10 mg/kg respectively) and received meloxicam (1 mg/kg, s.c.) for three consecutive days as postoperative analgesia. Viral expression and regional specificity were verified by immunohistochemistry, and mice showing inadequate expression were excluded from analysis.

### Neuro-Behavioural tests

#### Y-Maze

Spatio-temporal memory was assessed using a Y-maze with arms measuring 35 cm (length) × 7 cm (width) × 10 cm (height). During the exposure phase, mice were placed in the start arm and allowed to explore the start and familiar arms for 5 min, while the third arm was blocked. Animals were then returned to their home cages for a 10-min interval. In the test phase, mice were reintroduced to the start arm and allowed to explore the familiar and novel arms for 5 min. Time spent in each arm was recorded using ANY-maze behavioral tracking software (version 7). The maze floor was lined with corn bedding and replaced after each trial, while spatial cues were maintained throughout the experiment.

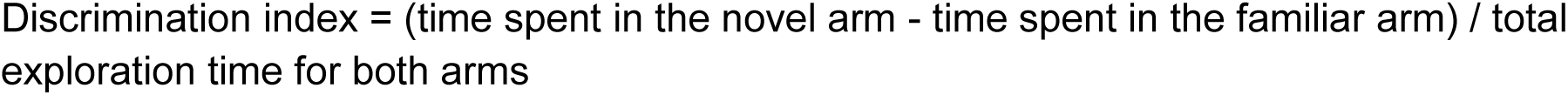

#### Social Recognition

Social recognition was assessed using a three-chamber apparatus (40 × 40 × 40 cm). Wire-mesh cages were placed in the two side chambers to hold stimulus mice. During habituation, the test mouse was placed in the center chamber and allowed to freely explore all chambers for 10 min. In the sociability phase, a stranger mouse was placed in a wire cage in one side chamber while the other cage remained empty, and the subject mouse was allowed to explore for 5 min. The mouse was then returned to its home cage for a 10-min interval. In the test phase, the familiar stranger mouse remained in one chamber and a second mouse from the subject’s home cage was placed in the opposite chamber. The subject mouse was allowed to freely explore, and exploration time (s) was recorded using ANY-maze behavioral tracking software (version 7). The discrimination index was calculated with the following equation:

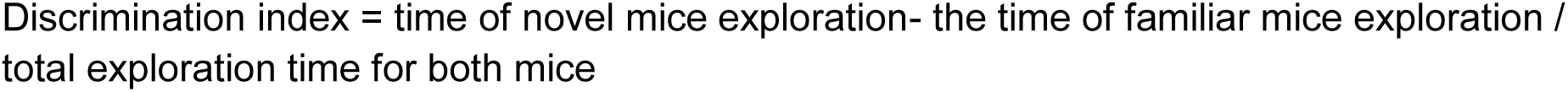

#### Open field test

The Open Field Test (OFT) was performed to assess locomotor and exploratory activity and to rule out nonspecific drug effects. Mice were placed in an open-field arena (45 × 45 × 40 cm) and allowed to freely explore for 60 min. After each session, animals were removed and the arena was cleaned with water and an odourless detergent to eliminate olfactory cues.

## RESULTS

### TGR5 is expressed in hippocampal Neural stem cells (_hi_NSC)

Considering impaired NSC homeostasis is strongly associated with aging associated neurological disorders (37), (38), and TGR5 activation attenuates neuroinflammation (39), we first determined TGR5 expression in cultured NSCs isolated from the hippocampi of postnatal day 0–1 mice pups. Immunocytochemical analysis revealed robust TGR5 immunoreactivity in NSCs **(Figure 1A)**. To determine whether this is consistent in the adult mice neurogenic niche, we analyzed mouse hippocampus using immunofluorescence and observed TGR5 immunoreactivity in SOX2+ NSCs in the sub granular region **(Figure 1B)**. To further determine the specificity of TGR5 expression, we performed lentiviral mediated knock-down of TGR5 in hippocampal NSCs and performed comparative Western blot and RT-PCR analysis for TGR5 expression. In both assays, we found significantly decreased expression of TGR5 with TGR5-shRNA in comparison to scrambled shRNA (scr-shRNA) transduced NSCs **(Figure 1 C, D)**. We also performed the immunofluorescence analysis of TGR5 expression in neurospheres expressing GFAP and nestin and observed the similar results **(Figure S1 A, B)**. We also found significant reduction in protein expression upon immunostaining of Scrambled and TGR5-ShRNA transduced neurospheres **(Figure S1 C, D)**. These *in vitro* and *in vivo* observation suggests that TGR5 is positioned to influence hippocampal NSC biology. This confirmed the specificity of TGR5 Expression over hippocampal Neural stem cell and to investigate the downstream functional role of TGR5 in neural stem cell physiology.

**Fig. 1.**
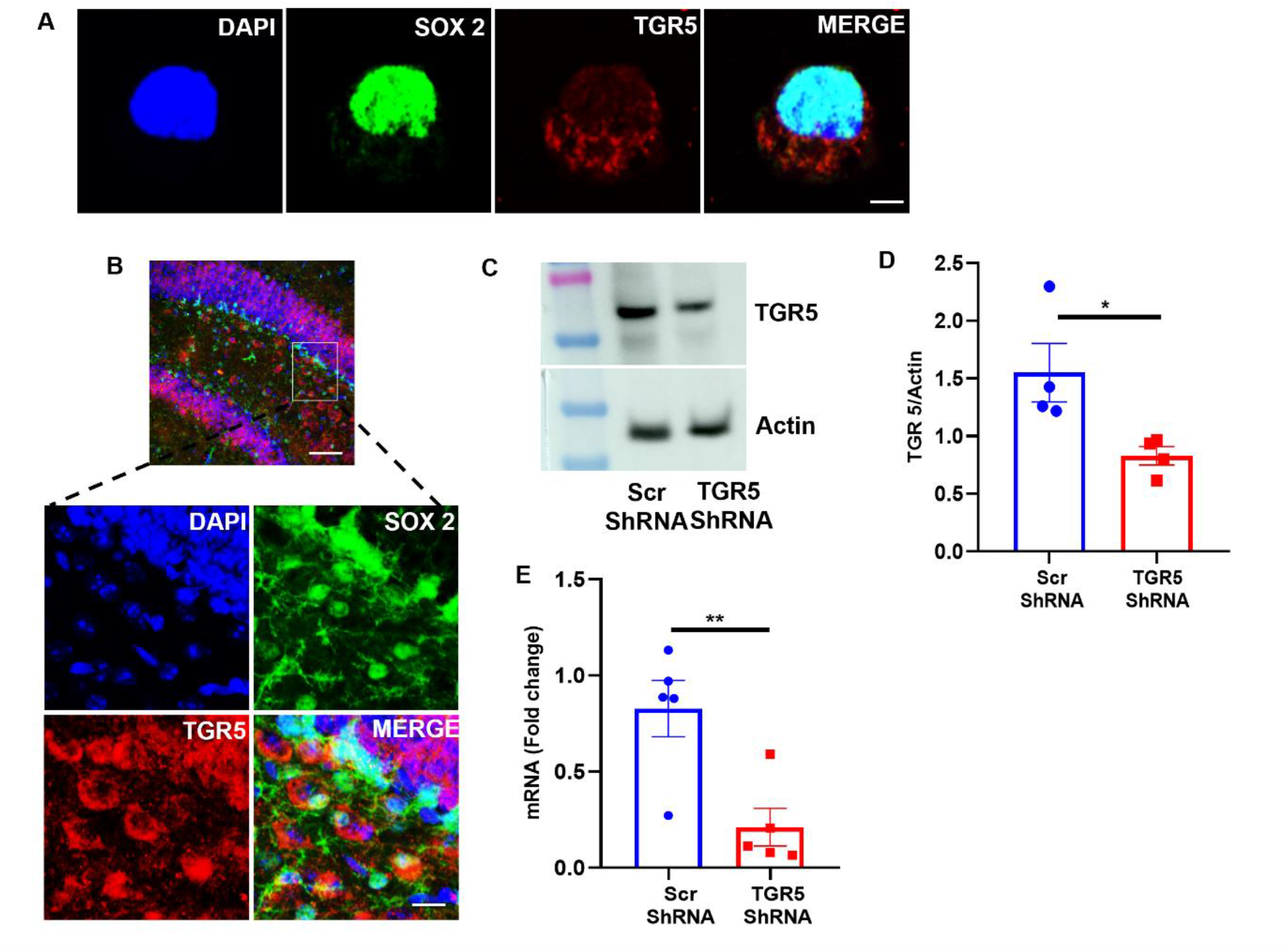
Expression of TGR5 in mice hippocampal neural stem cells (hiNSC). **(A)** Representative immunofluorescence image of _hi_NSC showing TGR5 expression in SOX2+ cells (Scale bar 2 µm). **(B)** Representative immunofluorescence image showing TGR5 expression in dentate gyrus of mice brain (scale bar 49 µm, 12 µm (magnified inset)). **(C)** Representative western blot of TGR5 expression in scrambled-shRNA (scr-shRNA) and TGR5-shRNA mediated knockdown in _hi_NSC. **(D)** Quantitation of western blots for TGR5 expression in _hi_NSC (C). Data were plotted as mean ± SEM, unpaired t-test, t = 2.719, *p = 0.0347, Scr-ShRNA vs TGR5-ShRNA n = 4 /group. **(E)** Quantitation of mRNA expression of TGR5 after Scr-shRNA or TGR5-shRNA mediated knockdown of TGR5. Data were plotted as mean ± SEM, unpaired t - test, t = 3.507, **p = 0.0080, Scr-ShRNA vs TGR5-ShRNA n = 5 /group.

### TGR5 induces NSC proliferation

Considering reduced NSC proliferative capacity in old aged brain (37), (38) and NSC activation may support endogenous neural repair, we first evaluated the effect of pharmacological activation of TGR5 on proliferation and differentiation of _hi_NSC. NSCs were treated with TGR5 agonist TC-G 1005 (1µM, 10µM, 72 h) and BrdU (10µM) followed by immunocytochemistry. Quantitative analysis of fluorescent images revealed that TC-G 1005 significantly increased the BrdU incorporation in concentration dependent manner, suggesting a proliferative role of TGR5 in NSCs **(Figure 2 A, B)**. To further support this observation, we determined the expression of PCNA in NSC lysate treated with TC-G 1005. Densitometric analyses of immunoblots revealed a concentration dependent increase in PCNA expression **(Figure 2, D)**. To determine the specificity of TC-G 1005 action, we performed the immunoblotting in Scr-shRNA or TGR5-shRNA transduced NSCs. As anticipated, we found TC-G 1005 treatment failed to induce any significant change in PCNA expression in TGR5-shRNA transduced NSCs in comparison to Scr-shRNA group **(Figure 2 E, F)**. To further investigate the effect of TGR5 agonist on NSC differentiation, we cultured NSC on PDL coated coverglass and maintained them in EGF/FGF free media and added TC-G 1005 (0.1µM, 1µM and 10µM) at every 72 h for 21 days followed by immunocytochemistry with nestin, DCX and MAP2. We found a concentration dependent increase in nestin expression, but decreased expression of MAP2, while no effect on DCX expression (**Figure S2 A, B, C, D**). These findings suggest that TGR5 agonism does not induce neuronal differentiation but instead promotes the stemness. These observations indicate that TGR5 activation modulates the self-renewal capacity of the _hi_NSC pool, preventing premature exhaustion (40).

**Fig. 2.**
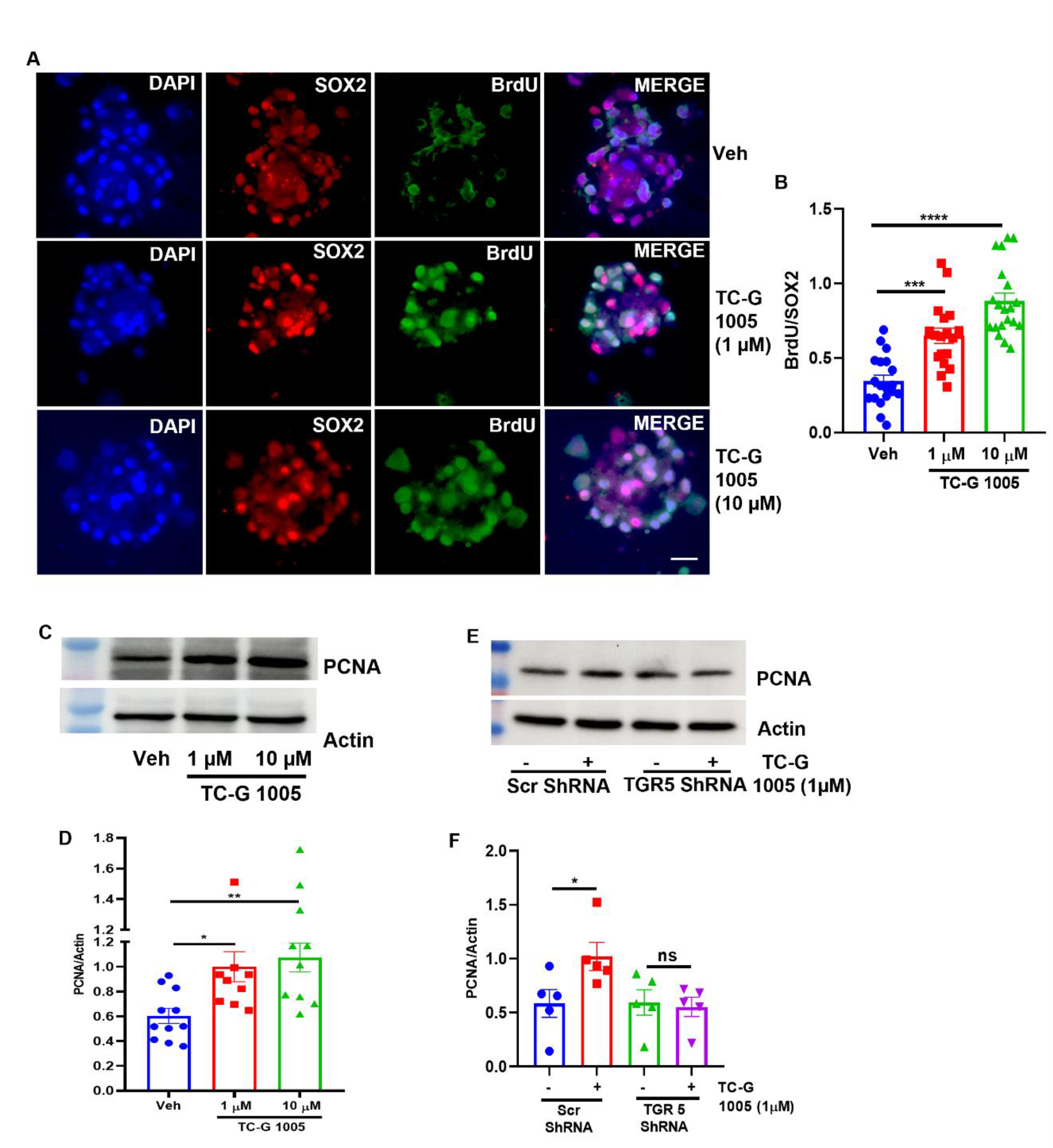
TGR5 activation stimulates _hi_NSC proliferation. **(A)** Representative images of dose dependent increase of BrdU incorporation in _hi_NSC upon TGR5 activation (Scale bar 25 µm). **(B)** Quantitation of BrdU incorporation in _hi_NSC as shown in (A). Data were plotted as mean ± SEM, Ordinary one-way ANOVA followed by Dunnett’s multiple comparisons test, (***p = 0.0001, Veh vs TC-G 1005 1µM), (****p < 0.0001 Veh vs TC-G 1005 10µM) n =3 independent experiment. **(C)** Representative western blot of PCNA showing its dose dependent increase in expression upon TGR5 activation. **(D)** Quantitation of PCNA shown in (C). Data were plotted as mean ± SEM, Ordinary one-way ANOVA followed by Dunnett’s multiple comparisons test, (*p = 0.0172, Veh vs TC-G 1005 1µM), (**p = 0.0047 Veh vs TC-G 1005 10µM) n =10 - 11 /group. **(E)** Representative western blot of PCNA showing no effect on proliferation upon TGR5 knockdown. **(F)** Quantitation of immunoblot shown in (E). Data were plotted as mean ± SEM, Ordinary one-way ANOVA followed by Sidak’s multiple comparisons test, (*p = 0.0372, Scr-ShRNA vs Scr-ShRNA + TC-G 1005 1µM, p = ns TGR5-ShRNA vs TGR5-ShRNA + TC-G 1005 1µM) n = 5 /group.

### TGR5 agonism modulate activation of quiescent neural stem cells and autophagy

NSC quiescence is essential homeostatic mechanism to preserve the NSC reservoir, and its dysregulation has been implicated in CNS disorders and age-related decline in neural plasticity (41), (16). Considering TGR5 agonisms promotes NSC proliferation while inhibiting differentiation into neurons, we sought to determine whether TGR5 activation could promote the transition of quiescent NSCs to an activated state. To test this, _hi_NSCs were exposed to BMP4 (50 ng/mL) for 4 days to induce quiescence (17), and were subsequently exposed with increasing concentrations of the TGR5 agonist TC-G 1005 (100nM –10µM; **Fig. 3A**). The effects of TC-G 1005 were assessed by measuring the expression of the quiescence and activation-associated protein FOXO3a (13), (14), Ascl1 (15), pEGFR (16) and proliferation marker PCNA. Western blotting analysis revealed that BMP4 significantly increased FOXO3a expression compared with vehicle treated cells, suggesting the induction of a quiescence in _hi_NSCs. Notably, TC-G 1005 significantly decreased FOXO3a expression compared with BMP4 alone, in concentration dependent manner **(Fig. 3 B, C)**. BMP4 also reduced PCNA expression compared with vehicle treatment, indicating suppression of proliferative activity _hi_NSCs **(Fig. 3 B, D)**. TC-G 1005 progressively restored PCNA expression, with the significant increase observed at 10µM, but effects observed at 100nM and 1µM were comparatively modest. Activated NSC are known to express phosphorylated EGFR (pY-1068) which is repressed in quiescent NSC. We found treatment with BMP4 significantly reduced expression of pEGFR, however total EGFR remain reduced in vehicle NSC depicting more of phosphorylation of EGFR maintaining stemness. Treatment with TC-G 1005 significantly increased the expression of pEGFR depicting activation of NSC from quiescent state **(Fig. 3 B, E)**. Thus, the reduction in FOXO3a and restoration of PCNA and pY1068 EGFR suggest that TGR5 activation shifts BMP4-treated _hi_NSCs away from quiescence and toward proliferative active state. Immunofluorescence analysis of Ascl1 expression in _hi_NSCs further confirmed that TGR5 activation promotes the reactivation of quiescent _hi_NSCs (**Fig. 3F**). Vehicle-treated cells exhibited SOX2 and Ascl1 immunoreactivity, whereas BMP4 markedly reduced Ascl1 staining while SOX2 expression remained detectable. These findings suggest that BMP4 suppressed the activated state without affecting stemness. Notably, the addition of 1 µM TC-G 1005 to BMP4-treated cultures restored Ascl1 immunoreactivity in SOX2-positive _hi_NSCs. Quantification of the Ascl1/SOX2 signal confirmed a significant reduction following BMP4 treatment and a significant recovery after TC-G 1005 exposure (**Fig. 3 G**). We also obtain similar trend upon western blotting of Ascl1 **(Fig. S3 A, B).**

**Fig. 3.**
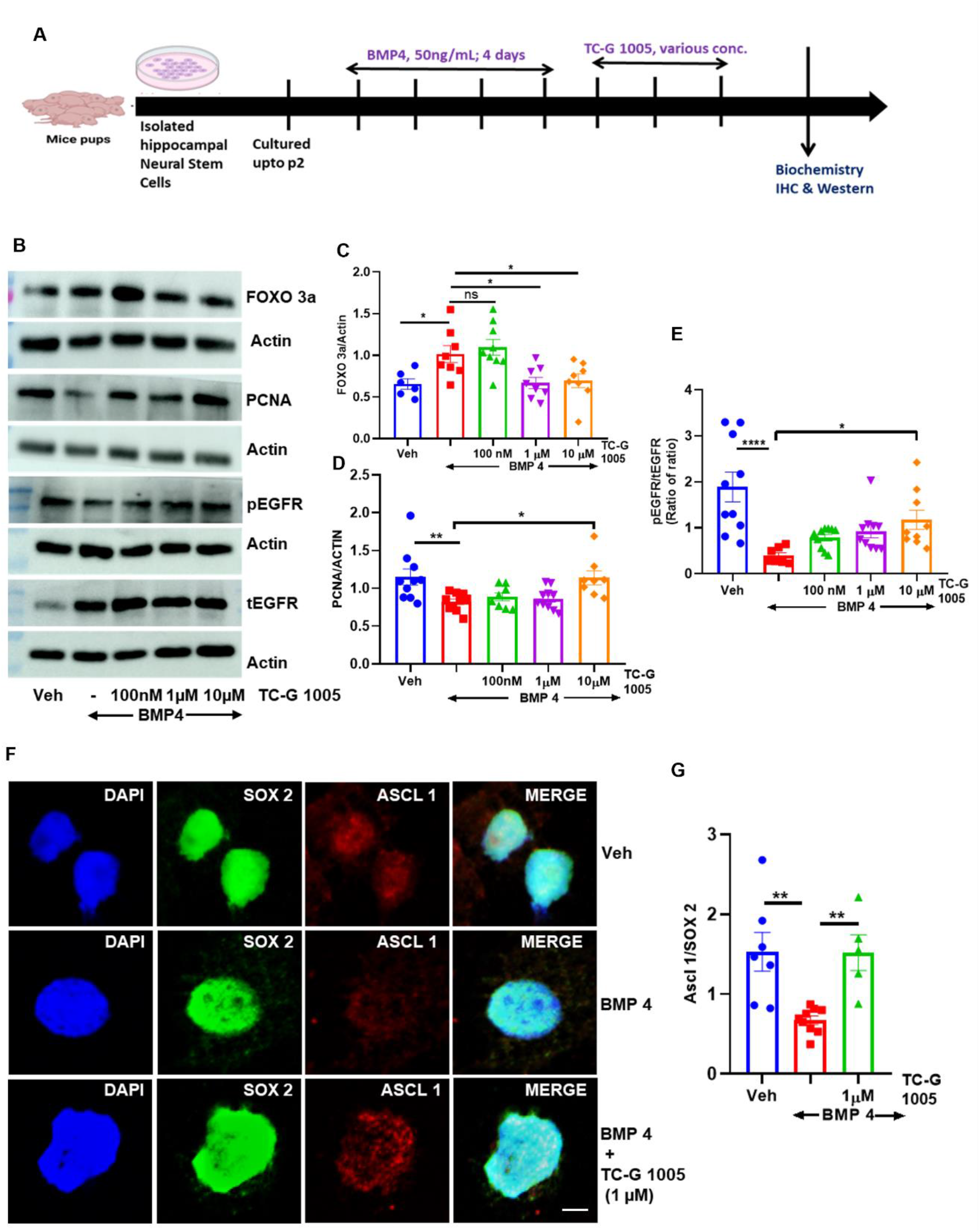
TGR5 activation reverse _hi_NSC quiescence. **(A)** Illustration of Quiescence establishment and TC-G 1005 treatment paradigm. **(B)** Representative western blots of FOXO3a, PCNA and total and phosphor EGFR in BMP4 & TC-G 1005 treated hiNSC showing dose dependent decrease in levels of FOXO3a & increase in levels of PCNA and Phospho EGFR depicting activation of hiNSC. **(C)** Quantitation of FOXO3a Expression as shown in (B). Data were plotted as mean ± SEM, Ordinary one-way ANOVA followed by Dunnett’s multiple comparisons test, (*p = 0.0308; Veh vs BMP4, p = ns BMP4 vs BMP4 + TC-G 1005 100 nM, *p = 0.0225; BMP4 vs BMP4 + TC-G 1005 1 µM, *p = 0.0394; BMP4 vs BMP4 + TC-G 1005 10 µM) n = 6 - 9 /group. **(D)** Quantitation of PCNA Expression as shown in (B). Data were plotted as mean ± SEM, Ordinary one-way ANOVA followed by Dunnett’s multiple comparisons test, (**p=0.0067; Veh vs BMP4, p = ns; BMP4 vs BMP4 + TC-G 1005 100 nM, p = ns; BMP4 vs BMP4 + TC-G 1005 1 µM, *p = 0.0148; BMP4 vs BMP4 + TC-G 1005 10 µM), n = 8 – 10 /group. **(E)** Quantitation of phospho EGFR and Total EGFR Expression as shown in (B). Data were plotted as mean ± SEM, Ordinary one-way ANOVA followed by Dunnett’s multiple comparisons test, (****p<0.0001; Veh vs BMP4, p = ns BMP4 vs BMP4 + TC-G 1005 100 nM, p = ns; BMP4 vs BMP4 + TC-G 1005 1 µM, *p = 0.0454; BMP4 vs BMP4 + TC-G 1005 10 µM), n = 7 - 10 /group. **(F)** Immunostaining of Ascl 1 in BMP4 & TC-G 1005 treated hiNSC showing its reappearance upon treatment (scale bar 8 µm). **(G)** Quantitation of expression of Ascl1 upon treatment as shown in (F). Data were plotted as mean ± SEM, Ordinary one-way ANOVA followed by Dunnett’s multiple comparisons test, (**p = 0.0028; Veh vs BMP4, **p = 0.0068 BMP4 vs BMP4 + TC-G 1005 1 µM) n = 3 independent experiment.

Since autophagy is intricately linked to NSC quiescence and activation (17), (18), we next examined whether TGR5 modulates autophagy during the transition of quiescent hiNSCs toward activation. Indeed, our western blot analysis showed that BMP4 exposure significantly increased p62 and LC3β-II levels, and reduced Beclin 1 expression **(Fig. 4 A)**, suggesting increased accumulation of autophagic machinery content in quiescent NSCs. In contrast, TC-G 1005 treatment following BMP4 exposure led to a dose-dependent reduction in p62 and LC3β-II levels and an increase in Beclin 1 expression, indicating the restoration of autophagic activity during NSC activation **(Fig. 4 B, C, D)**. Consistent with these findings, TC-G 1005 reduced LAMP2 puncta in _hi_NSCs (**Fig. 4 E, F**). Furthermore, analysis of _hi_NSC prepared from 0-1-day old mice pups of RFP-EGFP-LC3-tagged transgenic mice, BMP4 decreased the GFP signal, whereas TC-G 1005 increased it, suggesting enhanced lysosomal degradation **(Fig. S4 A, B)**. Collectively, these results demonstrate that pharmacological activation of TGR5 reverses key molecular features of BMP4-induced _hi_NSC quiescence, supporting the hypothesis that TGR5 agonism facilitates the transition of quiescent _hi_NSCs into an activated, proliferation-associated state.

**Fig. 4.**
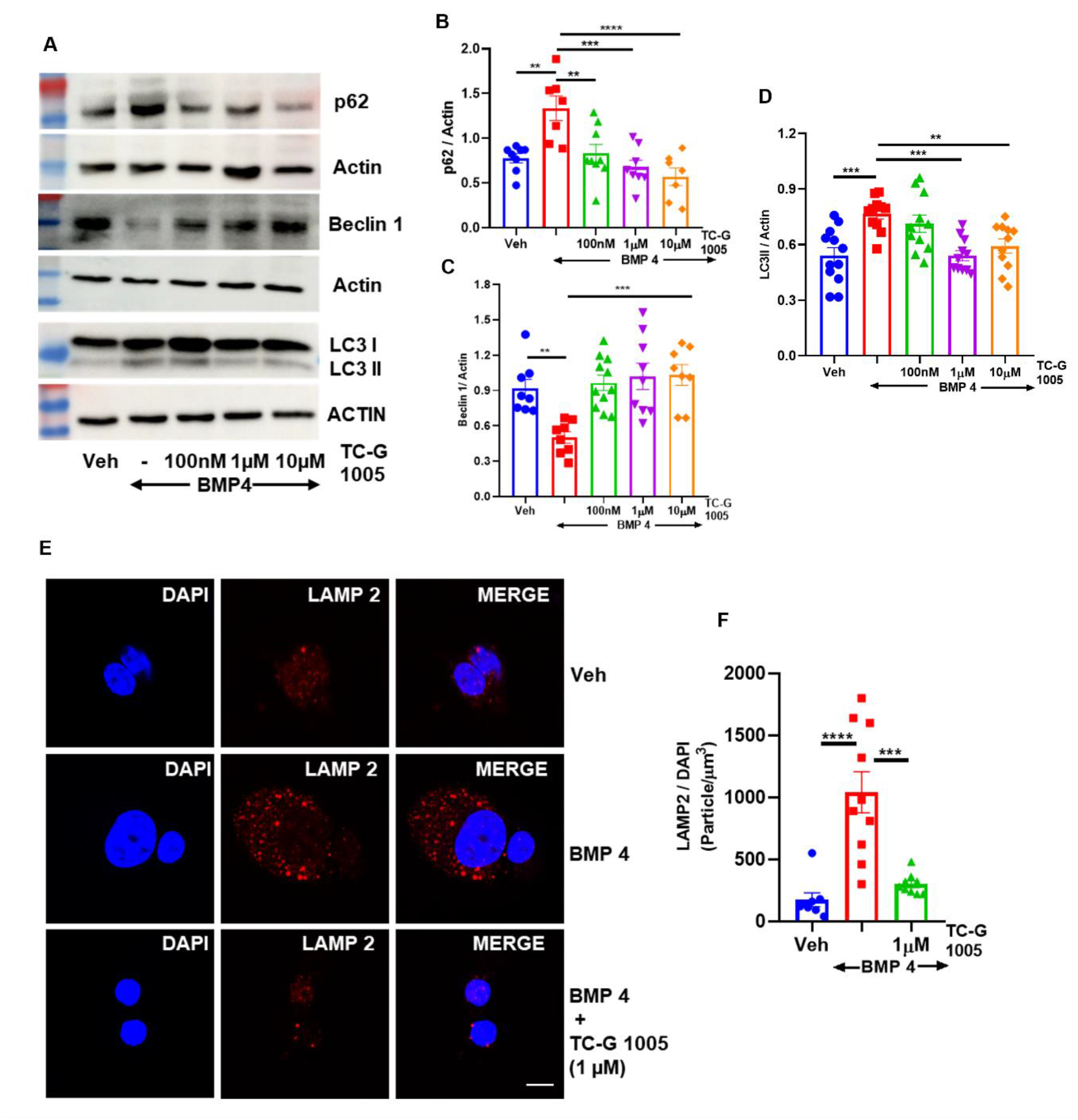
TGR5 modulates lysosome activity in qNSC and its transition to activation. **(A)** Representative western blots of p62, Beclin 1 & LC3 in BMP4 & TC-G 1005 treated hiNSC showing dose dependent decrease in levels of p62 & LC3-II, increase in beclin 1 depicting activation of hiNSC. **(B)** Quantitation of p62 expression as shown in (A). Data were plotted as mean ± SEM, Ordinary one-way ANOVA followed by Dunnett’s multiple comparisons test, (**p = 0.0011; Veh vs BMP4, **p = 0.0023; BMP4 vs BMP4 + TC-G 1005 100 nM, ***p = 0.0001; BMP4 vs BMP4 + TC-G 1005 1 µM, ****p < 0.0001; BMP4 vs BMP4 + TC-G 1005 10 µM) n = 7 - 9 /group. **(C)** Quantitation of Beclin1 expression as shown in (A). Data were plotted as mean ± SEM, Ordinary one-way ANOVA followed by Dunnett’s multiple comparisons test, (**p = 0.0050; Veh vs BMP4, ***p = 0.0007; BMP4 vs BMP4 + TC-G 1005 100 nM, ***p = 0.0003; BMP4 vs BMP4 + TC-G 1005 1 µM, ***p < 0.0003; BMP4 vs BMP4 + TC-G 1005 10 µM) n = 7 - 11 /group. **(D)** Quantitation of LC3 expression as shown in (A). Data were plotted as mean ± SEM, Ordinary one-way ANOVA followed by Dunnett’s multiple comparisons test, (***p = 0.0003; Veh vs BMP4, p = ns; BMP4 vs BMP4 + TC-G 1005 100 nM, ***p = 0.0004; BMP4 vs TC-G 1005 1 µM, **p = 0.0072; BMP4 vs BMP4 + TC-G 1005 10 µM) n = 12 - 13 /group. **(E)** Immunostaining of LAMP 2 in BMP4 & TC-G 1005 treated hiNSC showing its decrease upon treatment (scale bar 8 µm). **(F)** Quantitation of expression of LAMP2 upon treatment as shown in (E). Data were plotted as mean ± SEM, Ordinary one-way ANOVA followed by Dunnett’s multiple comparisons test, (****p < 0.0001; Veh vs BMP4, ***p = 0.0001 BMP4 vs BMP4 + TC-G 1005 1 µM) n = 3 independent experiment.

### Activation of TGR5 rescues neural stem cells from IL6 induced senescence

Chronic low-grade inflammation drives the transition from reversible quiescence to senescence (23), (22), (21). To determine whether TGR5 activation protects _hi_NSCs against inflammation-induced senescence, cultured _hi_NSCs were exposed to IL-6 (10 ng/mL, 24 h) and subsequently treated with increasing concentrations of the TGR5 agonist TC-G 1005 (100nM, 1µM, & 10µM) **(Fig. 5 A)**. Western blot analysis demonstrated that IL-6 markedly increased the expression of the senescence-associated proteins p53 and p16 compared with vehicle-treated cells **(Fig. 5 B, C, D)**. This increase was accompanied by a significant reduction in proliferating cell nuclear antigen (PCNA) **(Fig. 5 B, E)**, indicating that the acquisition of a senescence-associated phenotype was associated with impaired _hi_NSC proliferation. Furthermore, we also confirmed the effect of TC-G1005 on IL-6 induced _hi_NSC senescence by immunofluorescence and western blot analysis. We observed that IL-6 exposure markedly increased p16 staining within nestin-positive cell clusters, consistent with the induction of senescence in hiNSCs, while addition of TC-G 1005 after 24 h of IL-6 addition reduced p16 immunoreactivity, and preserved nestin expression. Quantification confirmed a significant reduction in the p16-to-nestin signal following TC-G 1005 treatment, with the strongest suppression observed at 10 µM (**Fig. 5 F, G**). We observed similar results with the western blot of p53 and p16 **(Fig. 5 B, C, D)**. These findings suggest that TGR5 signaling protects _hi_NSCs from inflammaging and promotes the recovery to the proliferative state. To further determine the specificity of TC-G 1005 action, we performed the western blotting and immunofluorescence of p16 in Scr-shRNA or TGR5-shRNA transduced NSC. As anticipated, we found TC-G 1005 treatment failed to induce any significant change in p16 expression of TGR5-shRNA transduced NSC in comparison to Scr-shRNA group **(Fig. S5 A, B, C, D).**

**Fig. 5.**
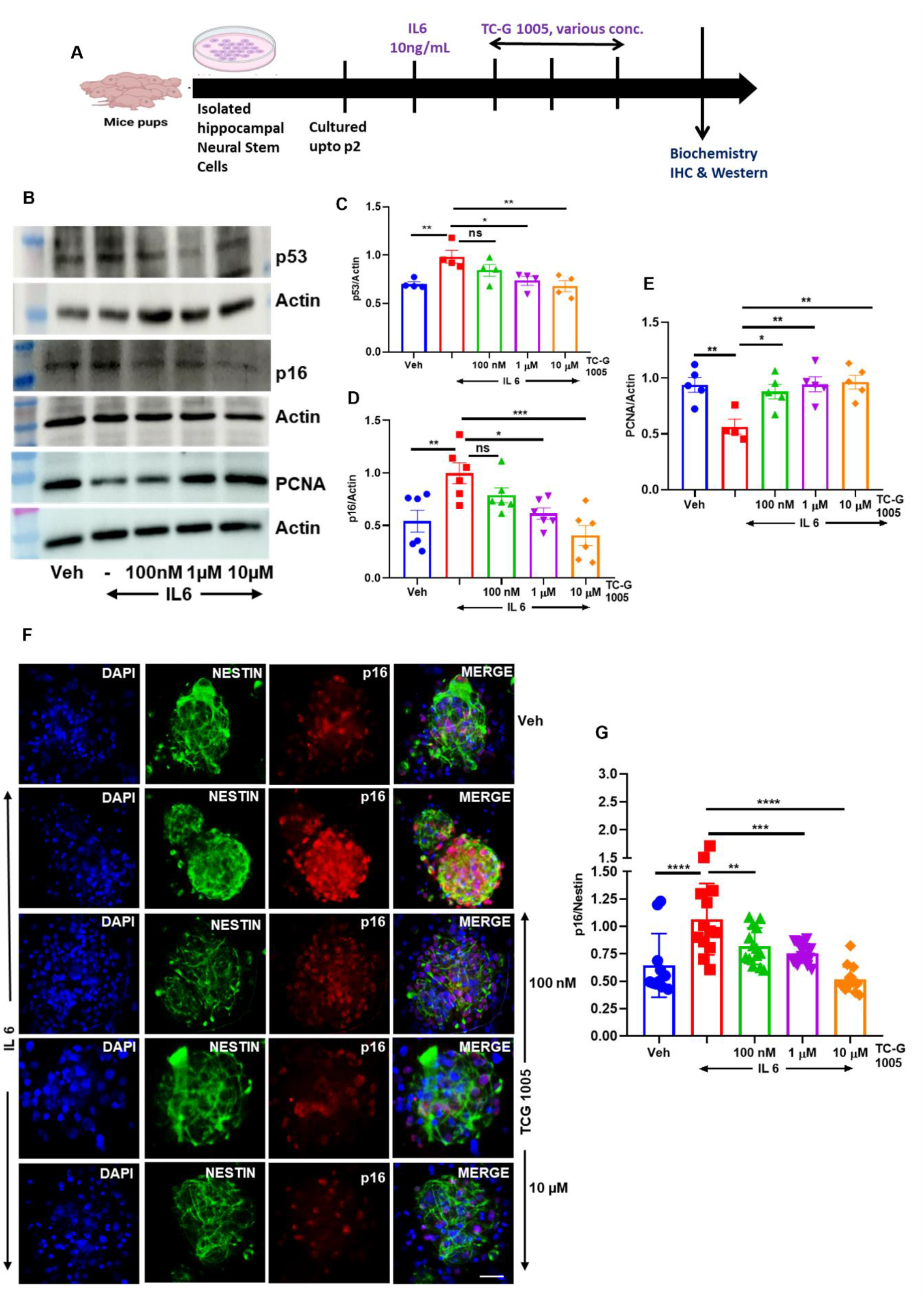
Activation of TGR5 rescues _hi_NSC from senescence. **(A)** Diagrammatic representation of IL6 induced inflammation and TC-G 1005 treatment paradigm. **(B)** Representative western blots of p53, p16 & PCNA in IL6 & TC-G 1005 treated _hi_NSC showing dose dependent decrease in levels of p53 & p16 depicting decrease in inflammation and increase in PCNA levels suggesting resumption of _hi_NSC proliferation and rescue from senescence. **(C)** Quantitation of p53 expression as shown in (B). Data were plotted as mean ± SEM, Ordinary one-way ANOVA followed by Dunnett’s multiple comparisons test, (**p = 0.0066; Veh vs IL6, p = ns; IL6 vs IL6 + TC-G 1005 100 nM, *p = 0.0156; IL6 vs IL6 + TC-G 1005 1 µM, **p = 0.0036; IL6 vs IL6 + TC-G 1005 10 µM) n = 4 /group. **(D)** Quantitation of p16 expression as shown in (B). Data were plotted as mean ± SEM, Ordinary one-way ANOVA followed by Dunnett’s multiple comparisons test, (**p = 0.0039; Veh vs IL6, p = ns; IL6 vs IL6 + TC-G 1005 100 nM, *p = 0.0159; IL6 vs IL6 + TC-G 1005 1 µM, ***p = 0.0002; IL6 vs IL6 + TC-G 1005 10 µM) n = 6 /group. **(E)** Quantitation of PCNA expression as shown in (B). Data were plotted as mean ± SEM, Ordinary one-way ANOVA followed by Dunnett’s multiple comparisons test, (**p = 0.0035; Veh vs IL6, *p = 0.0140; IL6 vs IL6 + TC-G 1005 100 nM, **p = 0.0032; IL6 vs IL6 + TC-G 1005 1 µM, **p = 0.0020; IL6 vs IL6 + TC-G 1005 10 µM) n = 4 -5 /group. **(F)** Immunostaining of p16 in IL6 & TC-G 1005 treated _hi_NSC showing its decrease upon treatment (scale bar 50 µm). **(G)** Quantitation of expression of p16 upon treatment as shown in (F). Data were plotted as mean ± SEM, Ordinary one-way ANOVA followed by Dunnett’s multiple comparisons test, (****p < 0.0001; Veh vs IL6, **p = 0.0095; IL6 vs IL6 + TC-G 1005 100 nM, ***p = 0.0003; IL6 vs IL6 + TC-G 1005 1 µM, ****p < 0.0001; IL6 vs IL6 + TC-G 1005 10 µM) n = 3 independent experiment.

### TC-G 1005 transcriptionally reverses IL-6-driven senescence in hippocampal NSCs

To elucidate the transcriptional programme underlying TC-G 1005 mediated rescue, we performed bulk RNA-sequencing on _hi_NSCs isolated from the hippocampi of 0-1-day old mouse pups and treated with vehicle, IL-6, or IL-6 + TC-G 1005 (n = 3 per group; **Fig. 6**), (treatment paradigm as in Fig. 5A). Treatment with IL-6 induced a marked transcriptional shift (660 genes upregulated and 82 genes downregulated Benjamini–Hochberg adjusted p ≤ 0.05, fold change > 1.5), consistent with a broad inflammatory reprogramming of the _hi_NSC pool **(Fig. 6 A)**. Addition of TC-G 1005 largely reversed this response, leaving only 49 differentially expressed genes relative to IL-6 alone (38 upregulated, 11 downregulated; **Fig. 6 B**), which is >90% contraction of the IL-6 signature that positions TGR5 agonism as a restorative rather than merely suppressive intervention. Principal component analysis resolved the three groups into discrete, minimally overlapping clusters, with the first two components capturing more than 85% of total variance, indicating that treatment was the dominant source of transcriptional variation rather than replicate or batch effects **(Fig. 6 C)**. Probing senescence-associated transcripts directly, TC-G 1005 reduced expression of the core cell-cycle-arrest effectors CDKN2a (p16^INK4a^), CDKN2b, CDK6, and the stress-response gene CALR (calreticulin), while increased the proliferation driving genes mTOR and HRAS **(Fig. 6 D)**. This bidirectional modulation of gene expression mirrors the recovery of proliferative capacity and loss of p16 immunoreactivity observed in vitro (Fig. 5B, E, F) and in the TGR5-activation experiments in vitro Fig. 2A. Furthermore, gene set enrichment analysis of the co-regulated transcripts also converged on NSCs maintenance and differentiation **(Fig. S6 A–D)**. The top 25 significant genes were similarly enriched for senescence and stemness regulators genes **(Fig. S6 E)**. These results collectively suggest that TGR5 activation transcriptionally uncouples IL-6 signalling from the senescence and quiescence programme, returning _hi_NSCs to a healthy and proliferative state.

**Fig. 6.**
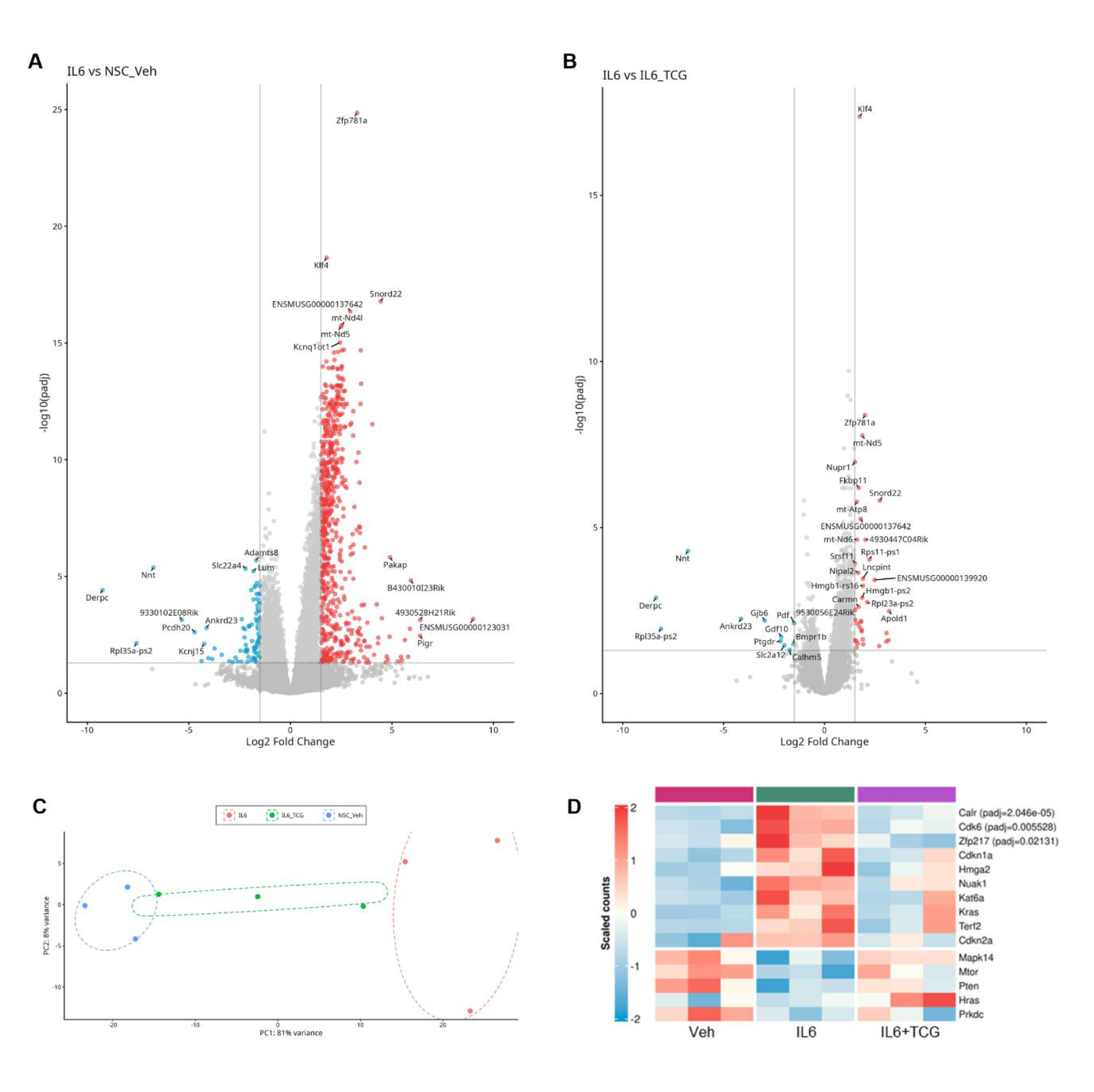
Bulk transcriptomic profile of _hi_NSCs. **(A, B)** Volcano plots of differentially expressed genes (DEGs) identified by DESeq2, showing IL-6 versus vehicle (A) and IL-6 versus IL-6 + TC-G 1005 (B). Red and blue denote significantly up- and down-regulated genes, respectively (Benjamini-Hochberg adjusted p ≤ 0.05); grey denotes non-significant genes. Vertical and horizontal lines mark fold-change and significance thresholds. **(C)** Principal component analysis of DESeq2-normalised counts. Each symbol represents one biological replicate, colour coded by treatment group; ellipses indicate group clustering. PC1 and PC2 together account for >85% of total variance. **(D)** Heatmap of row-scaled normalised counts for senescence- and proliferation-associated genes across vehicle, IL-6, and IL-6 + TC-G 1005 groups. Colour scale represents scaled counts (red-higher, blue-lower). Benjamini-Hochberg adjusted p-values are shown in parentheses for genes reaching p < 0.05.

### TGR5 agonist induces NSCs proliferation and suppress inflammation in aged mice

Given that chronic, low-grade sterile inflammation contributes to cognitive decline and increases the risk of dementia (42), (43), we examined whether systemic administration of the TGR5 agonist TC-G 1005 could reverse age-associated cognitive deficits and restore hippocampal NSC activity. Since the brain availability of TC-G1005 was not available, we performed mice plasma and brain pharmacokinetic analysis of TC-G 1005 to decide the optimal dosing in mice to determine its in vivo effects. As shown in supplementary table 1 and figure S7, we observed maximal concentration (C_max_) 224.71 ng/mL in plasma (equivalent to 0.56 µM) and 424.62ng/mL (equivalent to 1.06 µM) in brain. However, due to short half-life (0.68 h) of TC-G1005 in brain, we administered twice daily (5 mg/kg, s.c.) for 28 days to determine the effect of sustained activation of TGR5 on cognitive behaviours and NSCs homeostasis in18M old mice **(Fig. 7 A)**. As expected, aged mice exhibited a significantly lower Y-maze discrimination index than young mice, indicating impaired spatial or recognition memory. Chronic TC-G 1005 treatment significantly increased the Y-maze discrimination index in aged mice to the levels comparable to young mice **(Fig. S8 A)**. However, we didn’t observe any significant effect of TC-G 1005 treatment on social discrimination index **(Fig. S8 B)**. Open-field locomotor activity measurement showed that TC-G 1005 did not improve cumulative locomotor activity in old mice (**Fig. S8 C, D**), suggesting that the improvement in Y-maze performance was unlikely due to generalized locomotor stimulation.

**Fig. 7.**
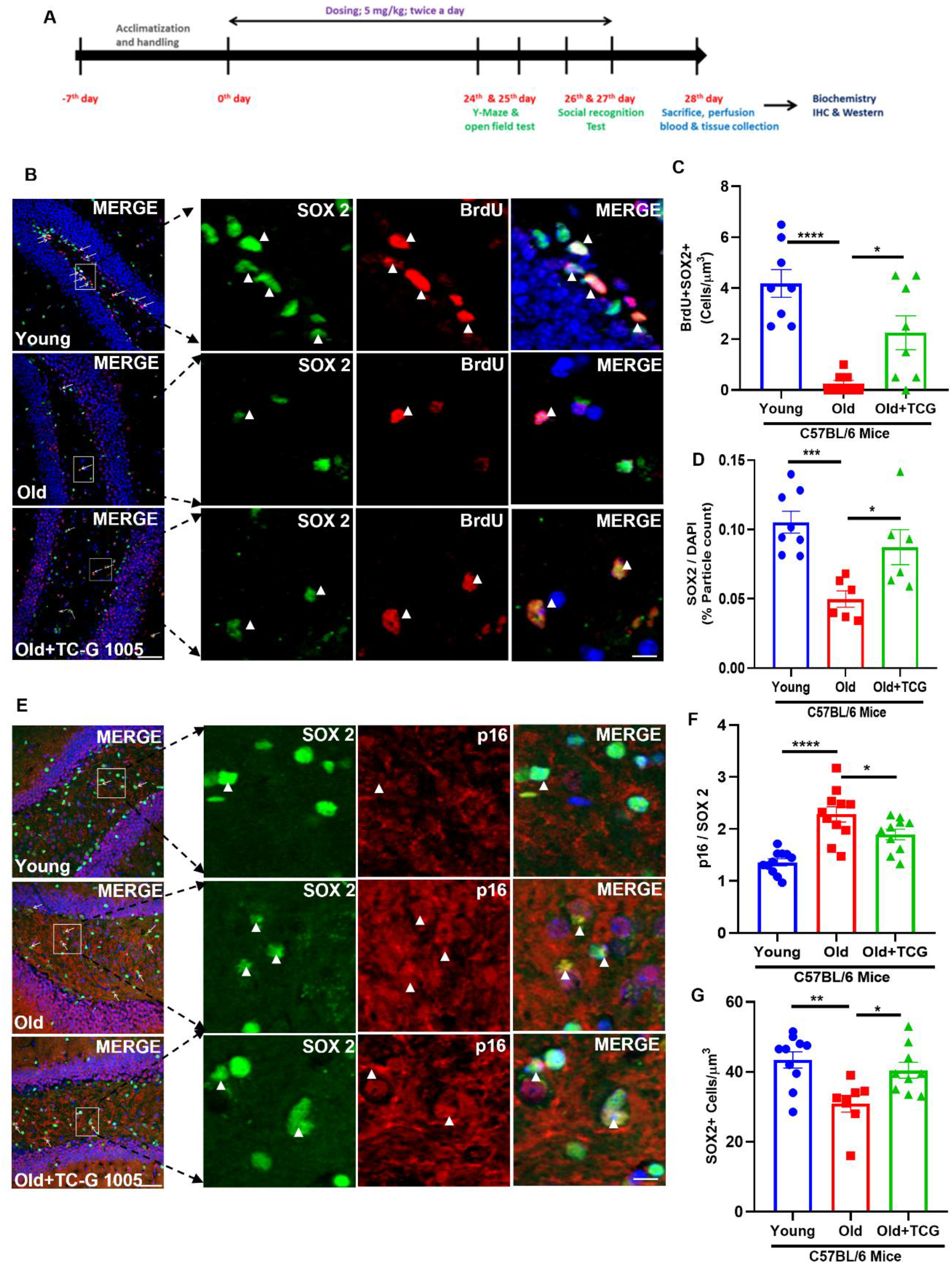
TGR5 augments cognitive impairment and senescence in aged mice. **(A)** Diagrammatic representation of drug treatment regimen and behavioral assay paradigm in 18-month-old C57BL6/J mice. **(B)** Representative images of dentate gyrus showing increase BrdU incorporation and increase in SOX2 population in old mice treated with TC-G 1005 suggesting proliferation of _hi_NSC and increase in stem cell pool (scale bar 49 µm). **(C)** Bar graph depicting quantitation of BrdU+SOX2+ cells as shown in (B). Data were plotted as mean ± SEM, Ordinary one-way ANOVA followed by Dunnett’s multiple comparisons test, (****p < 0.0001; Young vs Old, *p = 0.0194 Old vs Old + TCG 1005); n = 8/ group. **(D)** Bar graph depicting quantitation of SOX2+ cells as shown in (B). Data were plotted as mean ± SEM, Ordinary one-way ANOVA followed by Dunnett’s multiple comparisons test, (***p < 0.0008; Young vs Old, *p = 0.0240; Old vs Old + TCG 1005); n = 8/ group. **(E)** Representative images of dentate gyrus showing decrease in p16 expression in old mice treated with TC-G 1005 and increase in SOX2+ population (scale bar 49 µm). **(F)** Bar graph depicting quantitation of p16 as shown in (E). Data were plotted as mean ± SEM, Ordinary one-way ANOVA followed by Dunnett’s multiple comparisons test, (****p < 0.0001; Young vs Old, *p = 0.0398 Old vs Old + TCG 1005); n = 10-11/ group. **(G)** Bar graph depicting quantitation of SOX2+ cells as shown in (E). Data were plotted as mean ± SEM, Ordinary one-way ANOVA followed by Dunnett’s multiple comparisons test, (**p = 0.0019; Young vs Old, *p = 0.0186 Old vs Old + TCG 1005); n = 8-10/ group.

Importantly, immunofluorescence analysis of the hippocampal dentate gyrus of aged mice revealed a marked age-associated reduction in SOX2-positive neural stem/progenitor cells and proliferating BrdU/SOX2 double-positive cells. TC-G 1005 significantly increased the number of BrdU/SOX2 double-positive cells and the proportion of SOX2-positive cells in aged mice, indicating partial restoration of neural stem-cell proliferation **(Fig. 7 B, C, D)**. We also observed increased expression of p16 in SOX2-positive cells in aged mice and TC-G 1005 significantly reduced the p16/SOX2 signal, while increasing the overall number of SOX2-positive cells **(Fig. 7 E, F, G**). To assess the effect of TGR5 agonism on NSC quiescence, we analyzed the expression of Ascl 1 in SOX2+ cells in dentate gyrus region of mice. We found significant reduction in Ascl 1 expression levels in aged mice as compared to young mice, while TC-G treatment restored the number of Ascl 1+ NSCs. Moreover, we also found a total increase in number of SOX2+ cells in TC-G treated mice, suggesting activation of quiescent NSC to proliferative state **(Fig. S8 E, F, G)**. We also accessed the expression of DCX in aged mice DG region and found negligible DCX+ cells in aged and treated mice. Although interestingly, TGR5 Agonism increased the BrdU incorporation **(Fig. S8 H)**.

### Hippocampal TGR5 knockdown abrogated the TC-G 1005 effects

To investigate whether systemic TC-G 1005 requires hippocampal TGR5, we injected AAV9-TGR5-shRNA or AAV9-scrambled-shRNA (mCherry-tagged) bilaterally into the dentate gyrus of 18-month-old C57BL/6J mice, allowed three weeks of recovery, then dosed the animals with TC-G 1005 (5 mg/kg, twice daily, S.C.) for 28 days and tested behavior in the final week **(Fig. 8 A)**. Immunofluorescence staining for mCherry signal confirmed dentate gyrus-restricted viral targeting **(Fig. 8 B)**. The Y-maze test for spatiotemporal learning and memory showed that scrambled-shRNA injected mice discriminated the novel arm far more poorly than young mice (p = 0.0009). Notably, TC-G 1005 treatment in aged mice restored their discrimination index (p = 0.0140) in comparison to TGR5-shRNA injected aged mice **(Fig. 8 C)**. Social memory test also showed the same pattern **(Fig. 8 D)**. All aged groups travelled less than young mice in the open field, yet neither TC-G 1005 nor TGR5 knockdown changed total distance traveled **(Fig. 8 E)**, indicating locomotor activity is not associated with hippocampal TGR5 signaling. Knockdown also erased the cellular correlates of the drug effect. TC-G 1005 raised the density of BrdU⁺SOX2⁺ cells in the dentate gyrus of scrambled-shRNA mice (p = 0.0464), indicating that the resident neural stem cell pool resumed proliferation, but produced no such increase in TGR5-shRNA mice **(Fig. 8 F, G)**. In parallel, TC-G 1005 significantly reduced the number of senescent p16⁺SOX2⁺ NSCs in scrambled-shRNA mice (p = 0.0178) in comparison to TGR5-shRNA injected aged **(Fig. 8 H, I)**. These loss-of-function data identify hippocampal TGR5 as the necessary site of action for TC-G 1005. Losing both readouts together — stem cell proliferation and p16 clearance in TGR5 Knockdown group suggests hippocampal TGR5 knockdown abrogated the TC-G 1005 effects and TGR5 activation in scramble group restores cognition in the aged mice hippocampus.

**Fig. 8.**
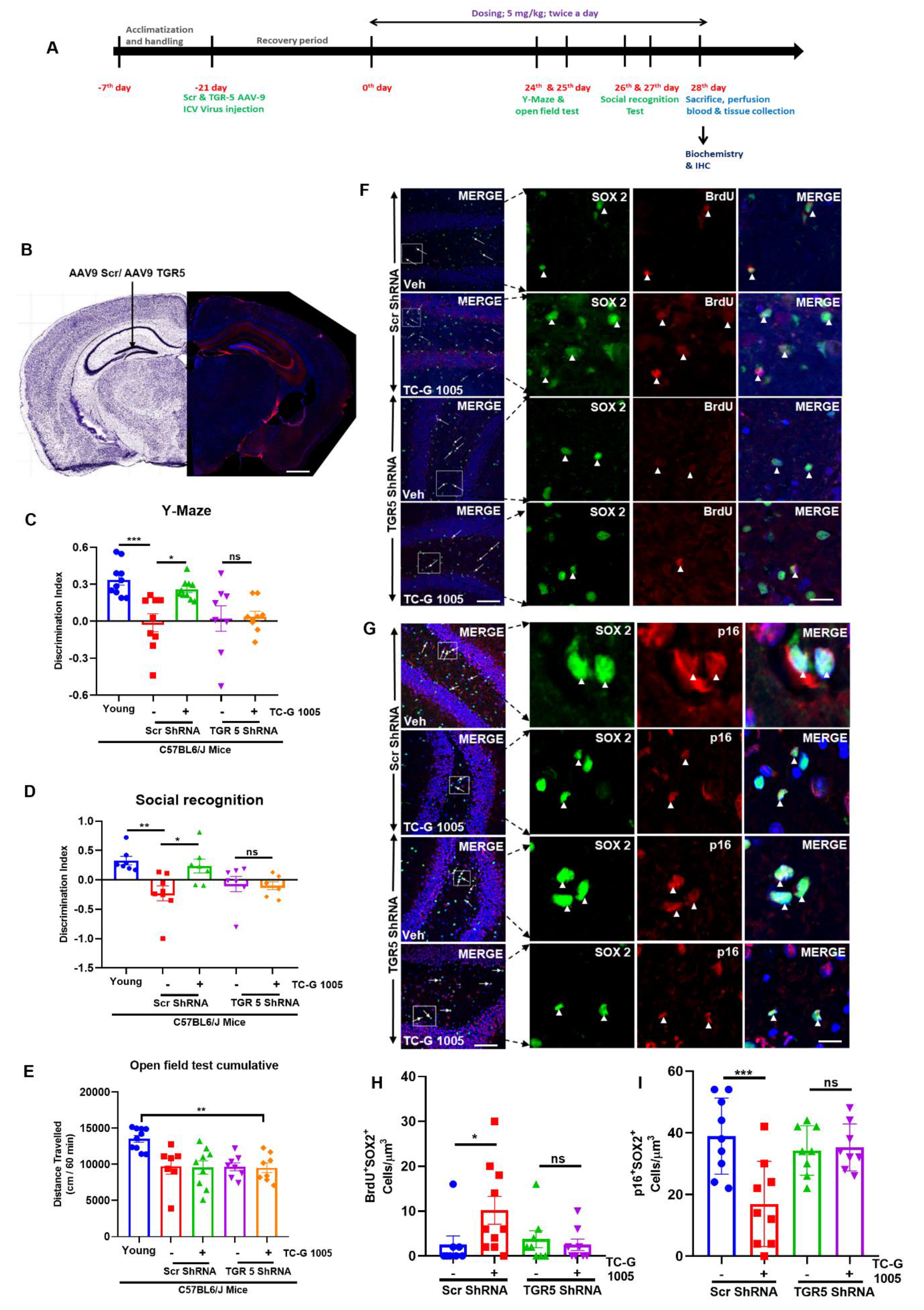
Hippocampal TGR5 knockdown abrogates TC-G 1005 induced NSCs proliferation, senescence & pro-cognitive effect of TC-G 1005 in aged mice. **(A)** Experimental timeline. **(B)** Representative images confirming dentate gyrus injection track by mCherry fluorescence (red) from AAV9-Scr-mCherry or AAV9-mTGR5-mCherry with DAPI (blue) (scale bar 2 inches). **(C)** Y-maze novel-arm discrimination index showing TC-G 1005 improved discrimination in Scr-shRNA aged mice but not in TGR5-shRNA aged mice. Data were plotted as Mean ± SEM; one-way ANOVA followed by Dunnett’s multiple comparisons test, ***p = 0.0009; young vs Scr-shRNA aged, *p = 0.0140 Scr-shRNA aged vs Scr-shRNA aged + TC-G 1005, p = ns TGR5-shRNA aged vs TGR5-shRNA aged + TC-G 1005; n = 8–10 mice/ group. **(D)** Social recognition discrimination index between a novel and a familiar conspecific mouse. TC-G 1005 restored social memory in Scr-shRNA mice. Data were plotted as Mean ± SEM; one-way ANOVA followed by Dunnett’s multiple test, **p = 0.0045 (young vs Scr-shRNA aged), *p = 0.0225 (Scr-shRNA aged vs Scr-shRNA aged + TC-G 1005), p = ns (TGR5-shRNA aged vs TGR5-shRNA aged + TC-G 1005); n = 7–8 mice/ group. (**E**) Cumulative distance travelled in the open field, recorded for 60 min in 5-min bins. All aged groups were hypoactive relative to young mice, but neither TC-G 1005 nor TGR5 knockdown altered locomotion. Data were plotted as Mean ± SEM; one-way ANOVA followed by Dunnett’s test, p = 0.0088, p = 0.0027, p = 0.0048 and p = 0.0035 for young versus Scr-shRNA aged, Scr-shRNA aged + TC-G 1005, TGR5-shRNA aged and TGR5-shRNA aged + TC-G 1005, respectively; ns = not significant, n = 7–10 mice / group. **(F)** Representative confocal images of the dentate gyrus immunostained for SOX2 (green), BrdU (red) and DAPI (blue) in Scr-shRNA and TGR5-shRNA aged mice treated with vehicle or TC-G 1005. Boxed regions are shown at higher magnification in the adjacent panels; arrowheads mark SOX2⁺BrdU⁺ double-positive cells (scale bar 49 µm). **(G)** Representative confocal images of the dentate gyrus immunostained for SOX2 (green), p16 (red) and DAPI (blue) across the same four groups, with magnified insets and arrowheads marking SOX2⁺p16⁺ senescent stem cells (scale bar 49 µm). **(H)** Quantification of BrdU⁺SOX2⁺ cell density from (F). TC-G 1005 increased proliferating NSCs in Scr-shRNA mice but not in TGR5-shRNA mice. Data were plotted as Mean ± SEM; one-way ANOVA with Dunnett’s multiple comparisons test, *p = 0.0464 (Scr-shRNA aged versus Scr-shRNA aged + TC-G 1005), p = ns (TGR5-shRNA aged versus TGR5-shRNA aged + TC-G 1005); n = 8–10 mice /group. **(I)** Quantification of p16⁺SOX2⁺ cell density from (H). TC-G 1005 reduced the senescent stem cell burden in Scr-shRNA mice, with no effect after TGR5 knockdown. Data were plotted as Mean ± SEM; one-way ANOVA with Sidak’s multiple comparisons test, *p = 0.0178 (Scr-shRNA aged versus Scr-shRNA aged + TC-G 1005), p = ns (TGR5-shRNA aged versus TGR5-shRNA aged + TC-G 1005); n = 6–8 mice/ group.

### Mouse brain proteomic changes during ageing and in response to TC-G 1005

Considering that TGR5 activation suppresses IL-6-induced neural stem cell (NSC) senescence, promotes NSC proliferation, and rescues BMP4-induced quiescence, we sought to determine whether pharmacological TGR5 agonism can similarly counteract ageing-associated proteomic remodelling in the hippocampal niche *in vivo*. Hippocampal lysates from young, aged, and TC-G 1005–treated aged mice were profiled by LC-MS/MS **(Fig 9 A)**, and unsupervised PLS-DA **(Fig 9 B)** revealed clear separation of young and aged proteomes along the first component, with treated-aged samples shifting away from the aged cluster toward an intermediate, partially rejuvenated position, an unbiased signature that TGR5 agonism measurably reprograms the aged hippocampal proteome. Further, the differential-abundance analysis (FC ≥ 1.2, adjusted p < 0.05) identified 271 ageing-altered proteins (O vs Y) and 238 treatment-responsive proteins (OT vs O), with 86 proteins shared between the two comparisons **(Fig 9 C)**, indicating that TC-G 1005 substantially engages the same protein network disrupted by ageing rather than acting through an orthogonal pathway. Volcano plots for each comparison **(Fig 9 D)** highlight the individual up- and down-regulated proteins, and a two-dimensional rescue-trajectory analysis plotting each protein’s ageing effect against its treatment effect **(Fig 9 Dc)** formally classifies the ageing-responsive proteome: 13 proteins were fully rescued and 7 partially rescued, 73 showed a directional trend toward reversal, 129 were treatment-specific, and 91 were unaffected. These clearly reflects that a meaningful fraction of the ageing signature is pharmacologically reversible. The top rescued proteins are resolved by group in a heatmap **(Fig 9 E)**, annotated by pathway membership and by ageing/inflammation/autophagy/oxidative-stress signatures, providing candidate mechanistic mediators. Further, pathway-enrichment analysis across the four protein sets **(Fig 9 F)** shows that TC-G 1005 preferentially restores metabolic, amino-acid, and cytosolic calcium-regulation pathways that are dysregulated with age, while several ageing-associated processes (e.g., lysosomal/vacuolar transport, complement- and oxidative-stress-related terms) remain only partially normalized, consistent with a treatment effect concentrated on metabolic and proteostatic rather than purely inflammatory axes. Finally, module-level analysis of four biologically curated protein sets **(Fig 9 G)** shows that TC-G 1005 significantly shifts the mitochondrial and complement C1q modules toward youthful levels (Mann-Whitney, individual proteins marked by fold change ≥ 1.2 and adjusted p < 0.05), while astroglial-reactivity and myelin proteins are less consistently normalized. These in vivo proteomic data analyses suggest that TGR5 agonism reverses a defined subset of ageing-associated hippocampal protein changes, particularly mitochondrial bioenergetic and complement/neuroinflammatory nodes, providing a molecular correlate for the anti-senescence and quiescence-rescuing activity of TGR5 signalling observed in cultured NSCs, and linking systemic TGR5 pharmacology to hippocampal stem-cell niche rejuvenation.

**Fig. 9.**
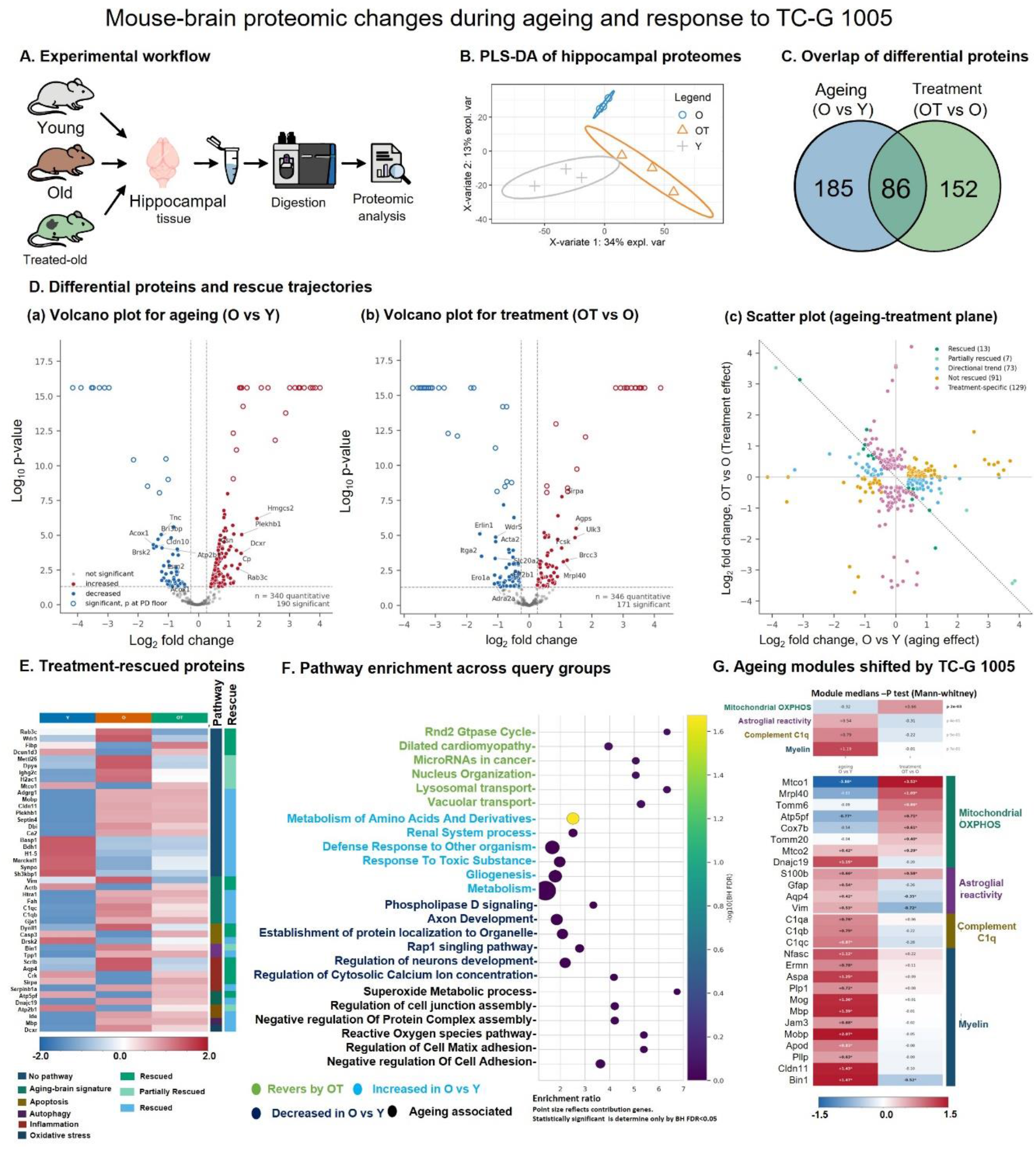
Global proteomic analysis of hippocampus of young (Y), 18M old (O) and old mice treated with TC-G 1005 (OT). **(A)** Experimental workflow: hippocampal tissue was collected from young, aged, and TC-G 1005-treated aged mice, digested, and analysed using LC-MS/MS. **(B)** Partial least-squares discriminant analysis (PLS-DA) of hippocampal proteomes, showing separation of young (Y), aged (O), and treated-aged (OT) groups along the first two components (34% and 13% explained variance, respectively). **(C)** Venn diagram of differentially abundant proteins (fold change ≥ 1.2, Proteome Discoverer-adjusted p < 0.05) in the ageing (O vs Y) and treatment (OT vs O) comparisons, showing 185 ageing-specific, 152 treatment-specific, and 86 shared differential proteins. **(D)** Differential proteins and rescue trajectories: (a, b) volcano plots for the ageing and treatment comparisons, respectively (red, increased abundance; blue, decreased abundance; grey, not significant; open circles, values at the Proteome Discoverer ratio-reporting floor/ceiling); selected proteins are labelled. Scatter plot of each protein’s ageing effect (log2 fold change, O vs Y) against its treatment effect (log2 fold change, OT vs O), classified as rescued (13), partially rescued (7), showing a directional trend (73), not rescued (91), or treatment-specific (129); the dotted line indicates exact reversal of the ageing effect. **(E)** Heatmap of the top 45 (of 93) treatment-rescued proteins, showing relative group profiles across Y, O, and OT, annotated by pathway association and rescue classification. **(F)** Pathway-enrichment analysis across four query groups shows that treatment with TC-G1005 reversed/restored pathways (green labels), increased in O vs Y (light blue), decreased in O vs Y (dark blue), and ageing-associated but not rescued (black) plotted by enrichment ratio, with point size reflecting the number of contributing genes and colour indicating −log10(BH-adjusted FDR). **(G)** Module-level analysis of four biologically defined protein sets (mitochondrial OXPHOS/import/mitoribosome, astroglia reactivity, complement C1q, myelin/oligodendrocyte) tested for a treatment effect against all other quantified proteins (Mann-Whitney); heatmap shows individual protein log2 fold changes for ageing and treatment, with asterisks marking |fold change| ≥ 1.2 and adjusted p < 0.05.

## Discussion

In this study we identified TGR5 as a pharmacologically tractable modulator of hippocampal NSC homeostasis and shown that its activation reverses two convergent features of the ageing niche, inflammation-driven senescence and deepened quiescence. TGR5 is expressed in SOX2⁺ NSCs in vitro and in the adult subgranular zone of DG, and its selective agonist TC-G 1005 increases BrdU incorporation and PCNA expression in a concentration-dependent and shRNA-sensitive manner. Together with the blockade of TGR5 agonist effect after hippocampus-restricted knockdown in aged mice, these data establish hippocampal TGR5 as both the molecular and anatomical site of action. Mechanistically, our data position TGR5 at the quiescence–activation switch rather than at the differentiation checkpoint. In BMP4-arrested NSCs, TC-G 1005 lowered FOXO3a, restored PCNA and pEGFR (Y1068), and recovered Ascl1 in SOX2⁺ cells, which are considered as the canonical signature genes for quiescence exit (13). This was further supported by normalization of autophagic flux as TGR5 agonism reduced p62 and LC3β-II, increased Beclin 1, and fewer LAMP2 puncta. Considering enhanced autophagolysosomal machinery content accumulation maintains NSC quiescence (18), (17), our findings suggest that TGR5-cAMP signalling relieves this lysosomal brake, a mechanistic bridge between bile acid sensing and stem cell state transitions. In parallel, TC-G 1005 suppressed IL-6 induced p53 and p16 expression and restored proliferation, and bulk RNA-sequencing showed that it tapered the IL-6 transcriptional signature by more than 90% while coordinately repressing Cdkn2a, Cdkn2b and Cdk6 and inducing mTOR and HRAS. The bidirectional nature of this shift argues that TGR5 agonism is restorative rather than simply anti-inflammatory.

Majority of above mentioned in vitro cellular effects translated in vivo as well. Twice-daily subcutaneous injection of TC-G 1005, a regimen justified by brain-to-plasma ratios (short brain half-life), restored Y-maze and social discrimination in 18-month-old mice without altering locomotion, expanded the SOX2⁺ and BrdU⁺SOX2⁺ pool, reduced p16⁺SOX2⁺ cells, and recovered Ascl1⁺ NSCs. Unbiased hippocampal proteomics showed that TC-G 1005 treated aged mice occupied an intermediate, partially rejuvenated position between young and aged proteomes, with 86 proteins shared between ageing- and treatment-responsive sets. Additionally, TC-G 1005 selectively normalised mitochondrial OXPHOS and complement C1q modules. Given that C1q accumulation and mitochondrial decline are among the most reproducible molecular hallmarks of brain ageing, the concordance between these modules and the anti-senescence phenotype provides a niche-level correlate for the cellular data and nominates candidate mediators for future investigations.

Despite increased NSC proliferation and reduced senescence burden, DCX⁺ immature neurons remained negligible in aged mice, and TC-G 1005 suppressed MAP2 while increased nestin in differentiating cultures. TGR5 activation therefore appears to replenish and de-repress the stem cell reservoir without forcing neuronal commitment, which is consistent with the gene set enrichment result favouring stem cell maintenance programmes. Cognitive benefit in treated aged mice may possibly arise due to decreased senescence-associated inflammatory and proteostatic burden within the niche rather than from net addition of new neurons, a mechanism that would circumvent the long-standing debate over adult human neurogenesis. However, this warrant further study to confirm whether the chronic activation of TGR5 induced expanded pool eventually yields BrdU⁺NeuN⁺ neurons and ablation of neurogenesis are necessary for the behavioural rescue. Sustained NSC proliferative stimulation by TGR5 agonism also raises the question of eventual pool exhaustion, which can be answered by longer dosing in future studies. Nonetheless, by linking a druggable bile acid receptor to NSC senescence and quiescence, and by demonstrating regional necessity, these findings identify TGR5 agonism as a candidate strategy for rejuvenating the aged neurogenic niche and preserving cognition.

## Authors

**Devanshu Kaushik:** Neuroscience and Ageing Biology Division, CSIR-Central Drug Research Institute, Lucknow, U.P. (226031), India. Jawaharlal Nehru University (JNU), New Delhi (110067), India.

**Prachi Dugam:** Neuroscience and Ageing Biology Division, CSIR-Central Drug Research Institute, Lucknow, U.P. (226031), India.

**Gourav Sarkar:** Molecular Metabolism & Cell Signaling Laboratory, Department of Biological Sciences & Bioengineering, Indian Institute of Technology Kanpur, UP 208016, India. Mehta Family Centre for Engineering in Medicine, Department of Biological Sciences & Bioengineering, Indian Institute of Technology Kanpur, UP 208016, India.

**Boda Arun Kumar:** Neuroscience and Ageing Biology Division, CSIR-Central Drug Research Institute, Lucknow, U.P. (226031), India. Jawaharlal Nehru University (JNU), New Delhi (110067), India.

**Sandip Kumar Patel:** Medical Research Council Toxicology Unit, University of Cambridge, Cambridge, UK.

**Sai Prasad Pydi:** Molecular Metabolism & Cell Signaling Laboratory, Department of Biological Sciences & Bioengineering, Indian Institute of Technology Kanpur, UP 208016, India. Mehta Family Centre for Engineering in Medicine, Department of Biological Sciences & Bioengineering, Indian Institute of Technology Kanpur, UP 208016, India. Gangwal School of Medical Sciences and Technology, Indian Institute of Technology Kanpur, UP, 208016, India.

## Authors Contribution

D.K. conducted the experiments, analysed and interpreted data, manuscript writing. P.D. helped in stereotaxic surgery and animal behaviour. G.S. analysed RNA sequencing data and generated figure 5 and S5. B.A.K prepared virus, S.K.P. Analysed proteomic data and generated figure 9. S.P.P helped in analysing RNA sequencing data and generating figure 5 and S5 P.N.Y. was in charge of the design of experiments, data interpretation, and manuscript writing.

## Funding Sources

This study was financially supported by the ICMR funded project GAP-0452 and CSIR funded project MLP-2034.

### Notes

The authors declared no conflict of interest.

## ACKNOWLEDGMENT

All the authors are thankful to the Director of CSIR-CDRI for providing the necessary research facilities. D.K. acknowledges the JNU and DBT, New Delhi, for research fellowships. The authors are also thankful to the SAIF and in-house Animal House Facility for behavioural studies

