## Supplementary information for "THE BILE ACID RECEPTOR TGR5 MODULATES HIPPOCAMPAL NEURAL STEM CELL STATE AND AGEING"

**Running Title:** Bile acid receptor and neural stem cell ageing (46 characters).

**Supplementary Table 1**. Pharmacokinetic profile of TC-G 1005 in C57BL6/j mice

| **TC-G 1005: Single Dose (10 mg/kg, S.C) Subcutaneous and Brain Distribution Study in Male C57BL/6 Mice** | | | |
| --- | --- | --- | --- |
| **Parameters** | **Mean Plasma PK Parameters** | **Parameters** | **Mean Brain PK Parameters** |
| Dose (mg/kg b.w.) | 10 | Dose (mg/kg b.w.) | 10 |
| C_max_ (ng/mL) | 224.71 | C_max_ (ng/g) | 424.62 |
| T_max_ (h) | 0.50 | T_max_ (h) | 0.50 |
| AUC_last_ (h*ng/mL) | 318.90 | AUC_last_ (h*ng/g) | 767.48 |
| AUC_inf_ (h*ng/mL) | 320.75 | AUC_inf_ (h*ng/g) | 781.20 |
| AUC_extrap_(%) | 0.58 | AUC_extrap_(%) | 1.76 |
| T_1/2_ (h) | 1.49 | T_1/2_ (h) | 0.68 |
| MRT_last_ (h) | 1.40 | MRT_last_ (h) | 0.94 |

MRT = Mean residence time; AUC = Area under curve; C_max_ = Maximum concentration; T_max_ = Maximum concentration time point; T_1/2_ = Half-life; b.w. = body weight

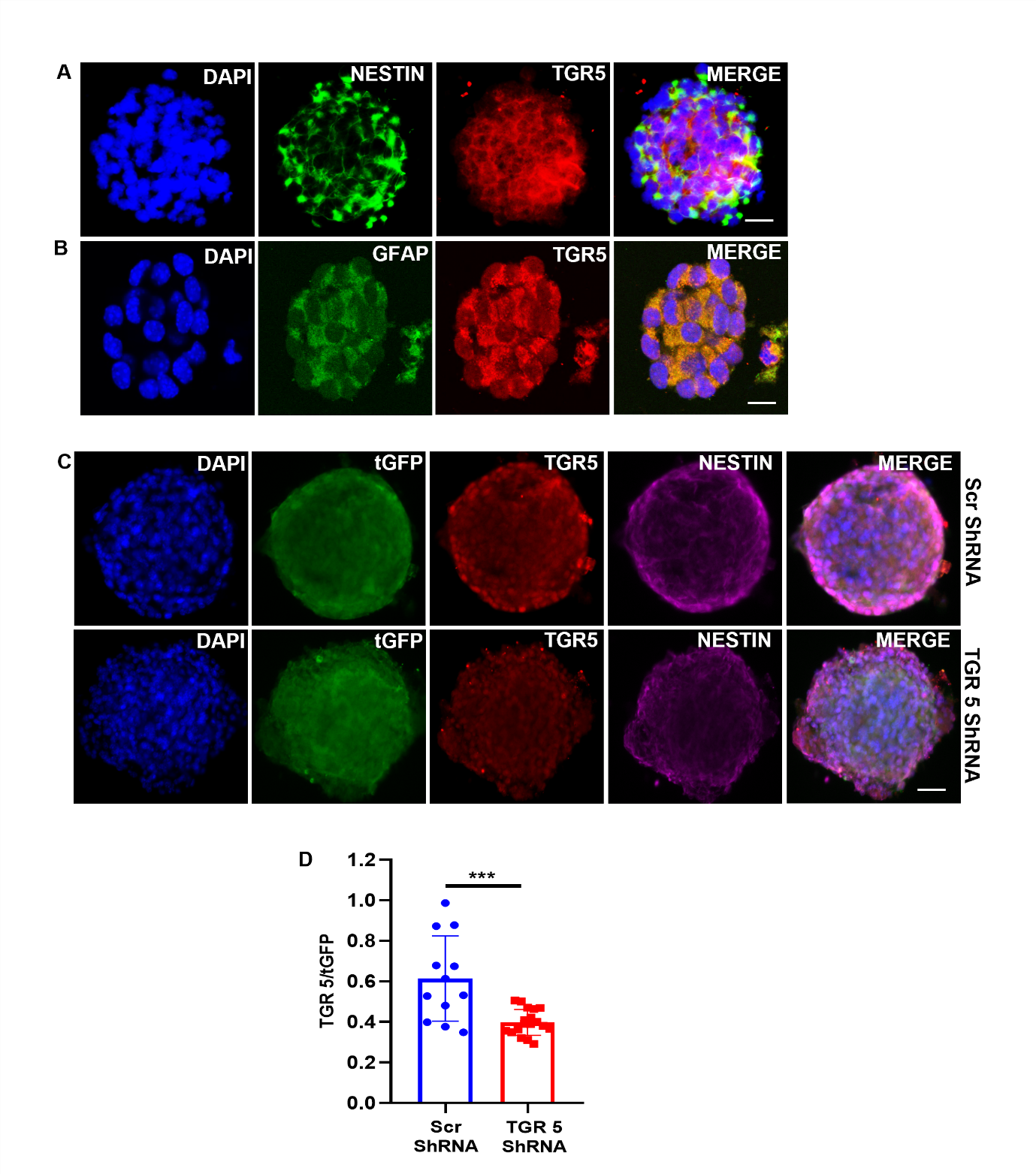

**Figure S1.** Expression of TGR5 in mice hippocampal neural stem cells (_hi_NSC). **(A & B)** Representative immunofluorescence image of _hi_NSC showing TGR5 expression in Nestin+ (Scale bar 50 µm) and GFAP+ (Scale bar 19 µm) neurosphere. **(C)** Representative immunofluorescent images of scrambled and TGR5 ShRNA transduced neurospheres depicting decrease in expression of TGR5 upon knockdown (Scale bar 50 µm). **(D)** Quantitation of TGR5 expression upon lentiviral mediated knockdown as shown in (C). Data were plotted as mean± SEM, unpaired t-test, t = 4.113, ***p = 0.0003, Scr-ShRNA vs TGR5-ShRNA n = 3 independent experiment.

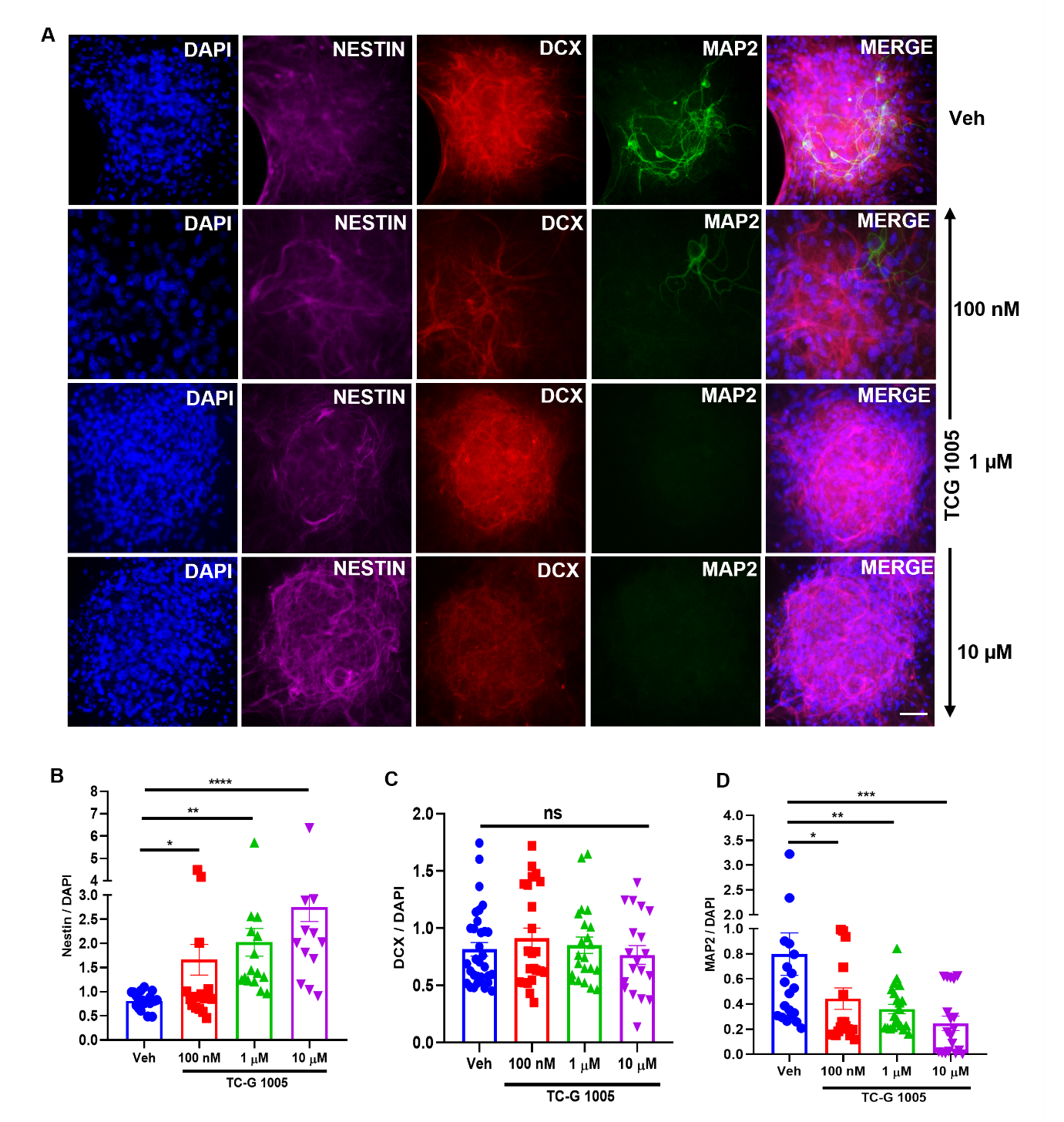

**Figure S2. (A)** Representative images showing expression nestin (marker of NSC), DCX (marker of immature neuron) and MAP2 (Marker of mature neuron) in neurospheres treated with various concentration of TC-G 1005 at every 72 h for 21 days. (Scale bar 50 µm)**.** **(B)** Quantitation of nestin as shown in (A) depicting dose dependent increase in expression of nestin. Data were plotted as mean ± SEM, Ordinary one-way ANOVA followed by Dunnett’s multiple comparisons test, (*p = 0.0486; Veh vs TC-G 1005 100 nM, **p = 0.0038; Veh vs TC-G 1005 1 µM, ****p<0.0001; Veh vs TC-G 1005 10 µM, n = 3 independent experiment. **(C)** Quantitation of DCX as shown in (A) depicting no effect on DCX. Data were plotted as mean ± SEM, Ordinary one-way ANOVA followed by Dunnett’s multiple comparisons test, (p = ns; Veh vs TC-G 1005 100 nM, p = ns; Veh vs TC-G 1005 1 µM, p = ns; Veh vs TC-G 1005 10 µM, n =3 independent experiment. **(D)** Quantitation of MAP2 as shown in (A) depicting dose dependent decrease in Expression of MAP2. Data were plotted as mean ± SEM, Ordinary one-way ANOVA followed by Dunnett’s multiple comparisons test, (*p = 0.0420; Veh vs TC-G 1005 100 nM, **p = 0.0072; Veh vs TC-G 1005 1 µM, ***p < 0.0009; Veh vs TC-G 1005 10 µM, n =3 independent experiment.

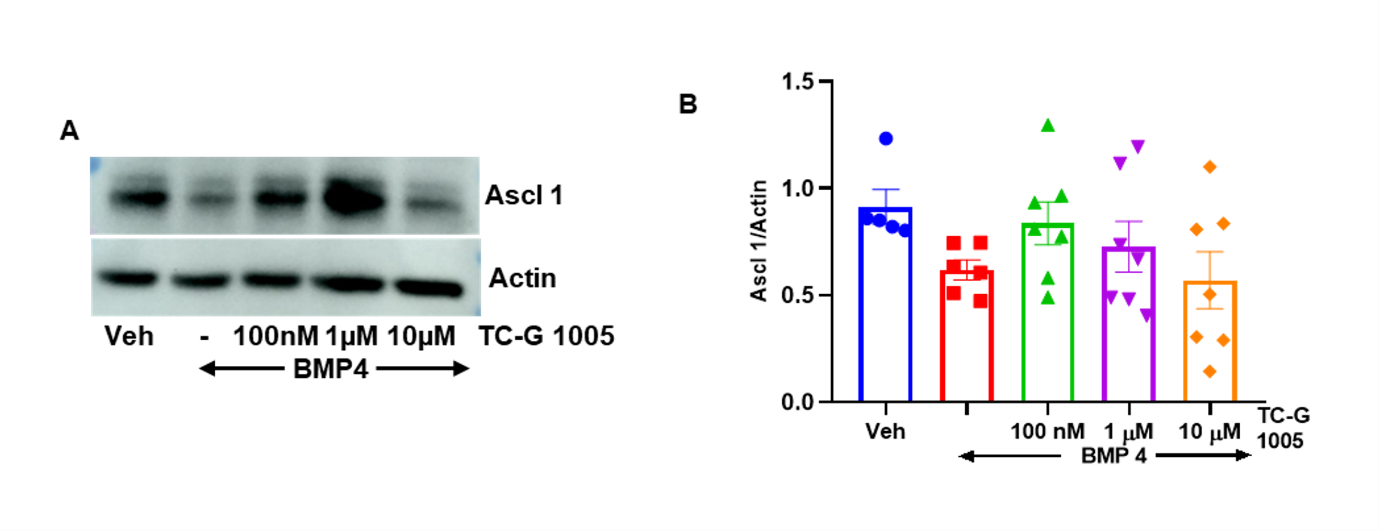

**Figure S3. (A)** Representative western blots of Ascl 1 in BMP4 & TC-G 1005 treated _hi_NSC showing dose dependent increase in levels of Ascl 1 depicting activation of _hi_NSC. **(B)** Quantitation of Ascl 1 Expression as shown in (A) showing a possible trend.

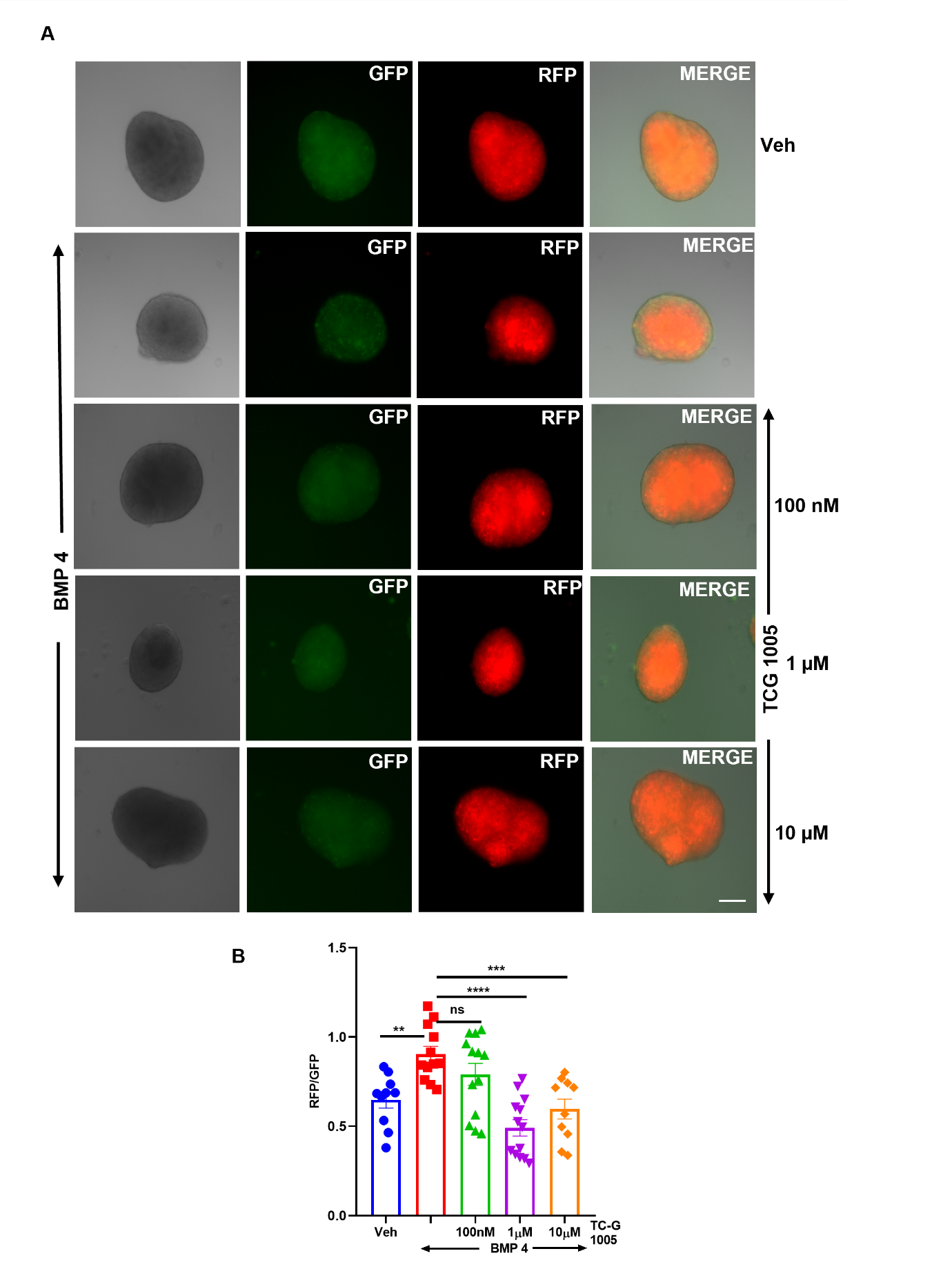

**Figure S4. (A)** Live imaging of hiNSC culture from RFP-GFP-LC3 mice pups treated with BMP4 and TC-G 1005 showing decrease in expression of GFP in qNSC depicting presence of numerous large autolysosomes which are not yet degraded in qNSC and increase in expression of GFP upon treatment with TC-G 1005 suggesting resumption of autophagy and transition to aNSC. (Scale bar 250 µm). **(B)** Quantitation of RFP to GFP Ratio as shown in (A). Data were plotted as mean ± SEM, Ordinary one-way ANOVA followed by Dunnett’s multiple comparisons test, (**p = 0.0048; Veh vs BMP4, p = ns; BMP4 vs BMP4 + TC-G 1005 100 nM, ****p < 0.0001; BMP4 vs BMP4 + TC-G 1005 1 µM, ***p = 0.0006; BMP4 vs BMP4 + TC-G 1005 10 µM) n = 3 independent experiment.

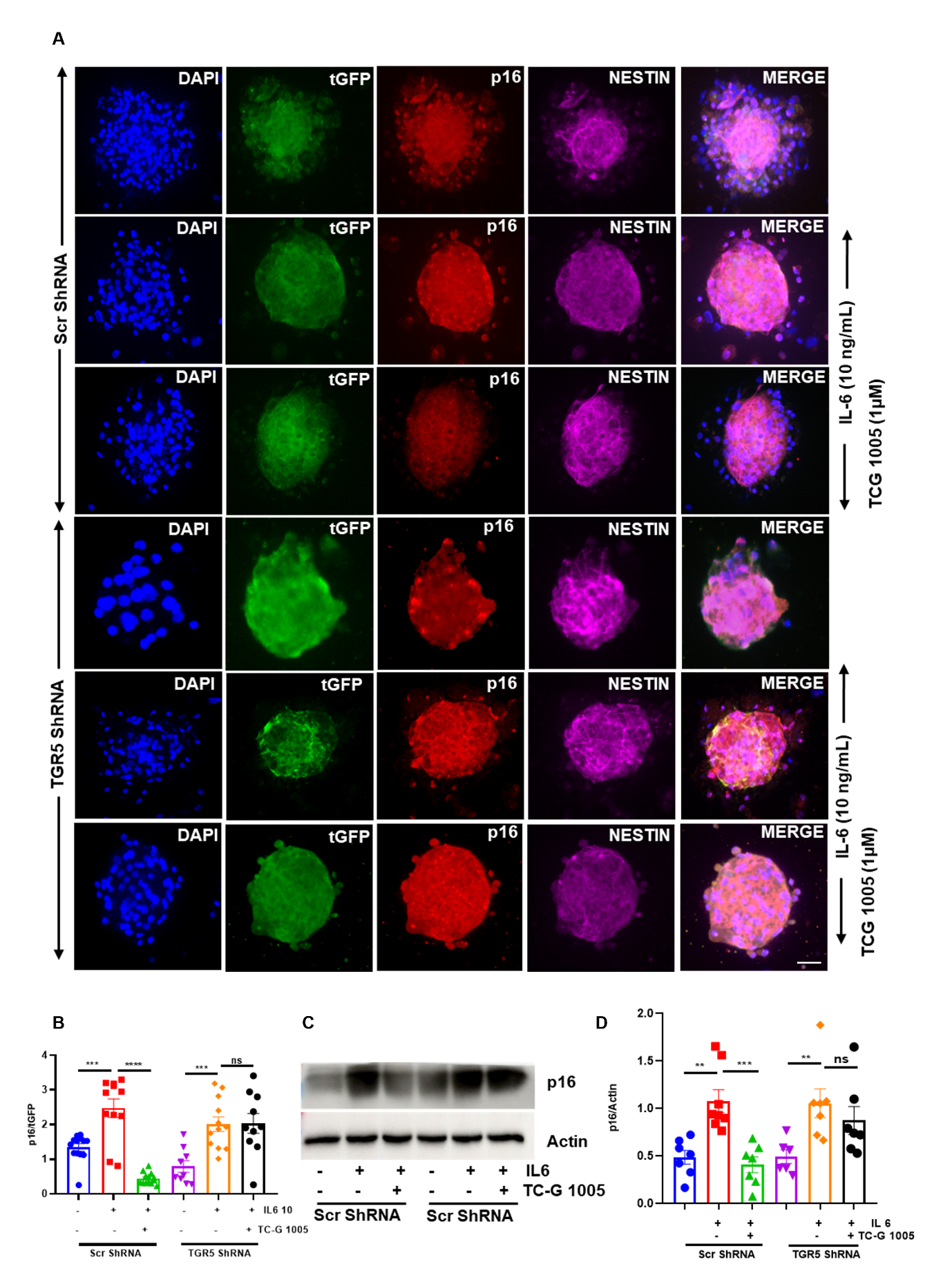

**Figure S5. (A)** Immunostaining of p16 in neurosphere with lentiviral transduced scrambled and TGR5 ShRNA treated with IL6 & TC-G 1005 treated hiNSC showing no effect of TC-G 1005 upon TGR5 knockdown. (Scale bar 50 µm). **(B)** Quantitation of p16 expression as shown in (A). Data were plotted as mean ± SEM, Ordinary one-way ANOVA followed by Sidak’s multiple comparisons test, (***p = 0.0002; Scr-ShRNA vs Scr-ShRNA + IL6, ****p < 0.0001 ; Scr-ShRNA + IL6 vs Scr-ShRNA + IL6 + TC-G 1005 1 µM, ***p = 0.0002; TGR5-ShRNA vs TGR5-ShRNA + IL6, p = ns; TGR5-ShRNA vs TGR5-ShRNA + IL6 + TC-G 1005 1µM) n = 3 independent experiment. **(C)** Representative western blot of p16 in lentiviral transduced scrambled and TGR5 ShRNA in IL6 & TC-G 1005 treated _hi_NSC showing no effect of TC-G 1005 upon TGR5 knockdown. (D) Quantitation of expression of p16 upon treatment as shown in **(D)**. Data were plotted as mean ± SEM, Ordinary one-way ANOVA followed by Sidak’s multiple comparisons test, (**p = 0.0023; Scr-ShRNA vs Scr-ShRNA + IL6, ***p < 0.0005 ; Scr-ShRNA + IL6 vs Scr-ShRNA + IL6 + TC-G 1005 1 µM, **p = 0.0083; TGR5-ShRNA vs TGR5-ShRNA + IL6, p = ns; TGR5-ShRNA vs TGR5-ShRNA + IL6 + TC-G 1005 1µM) n = 6 - 8 independent experiment.

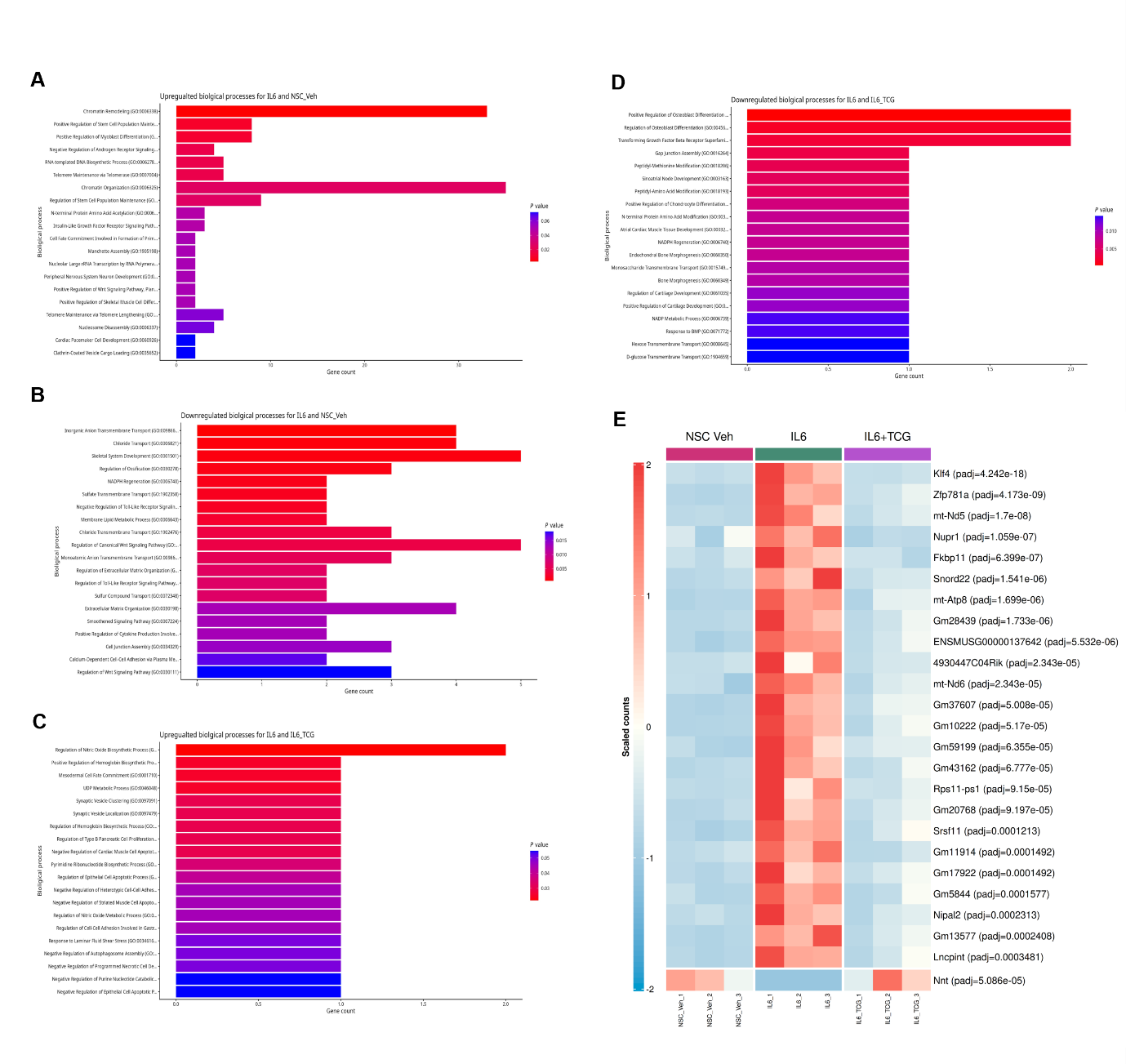

**Figure S6.** Gene Ontology (GO) analysis of DEGs in **(A & B)** Vehicle vs IL6 Treated **(C & D)** and IL6 Treated vs IL6 + TC-G 1005 Treated DESeq2 comparisons. **(E)** Heatmap showing the normalised counts for top most 25 significant DEGs in IL6 compared to IL6 with TCG treatment. The Benjamini Hochberg-adjusted *p*-value for each gene is shown in parenthesis if below 0.05 for panel (E).

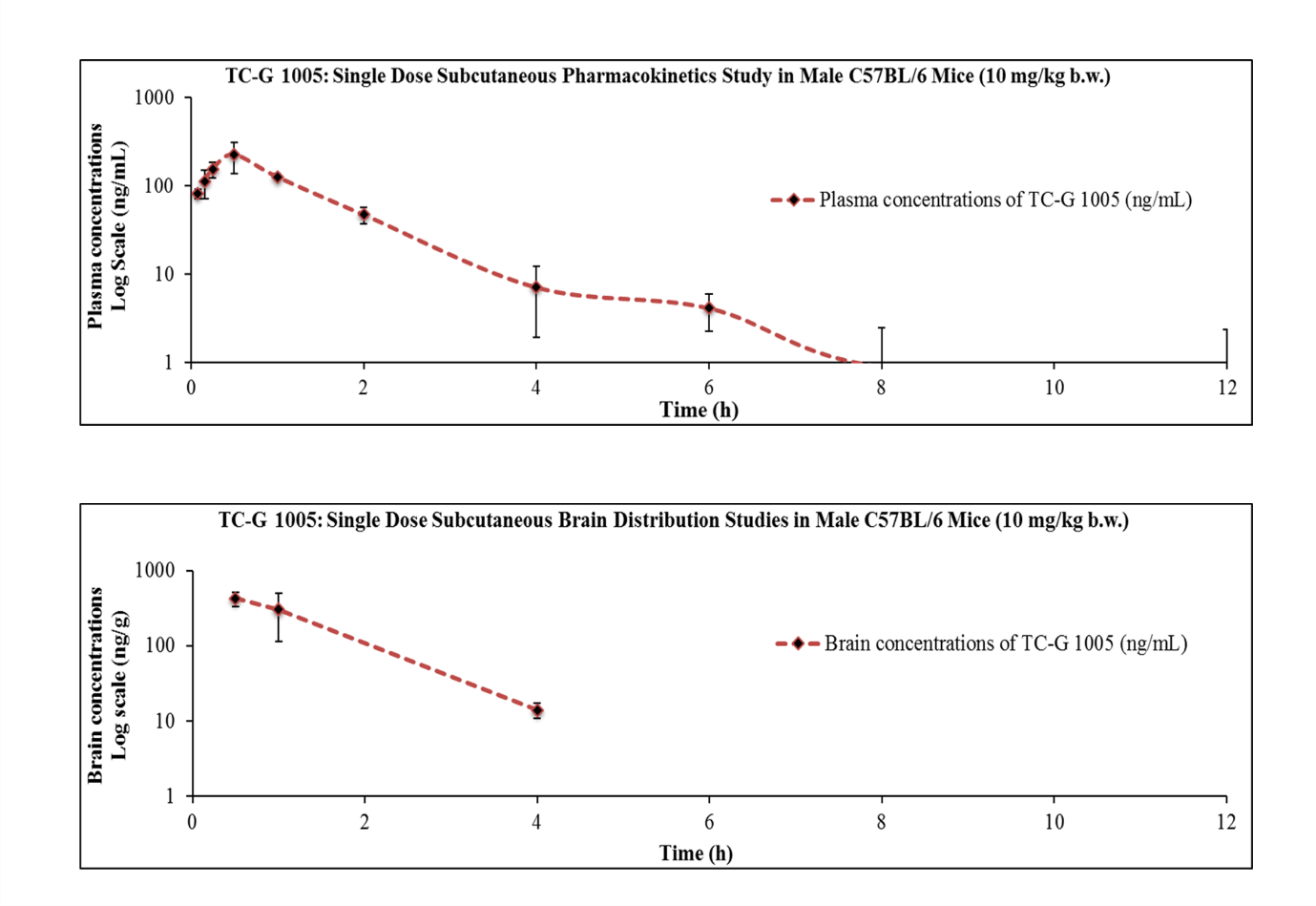

**Figure S7.** Pharmacokinetic profile of TC-G 1005 in plasma and brain tissue after Subcutaneous administration at 10 mg/kg in male C57BL/6j mice (n = 3/time point).

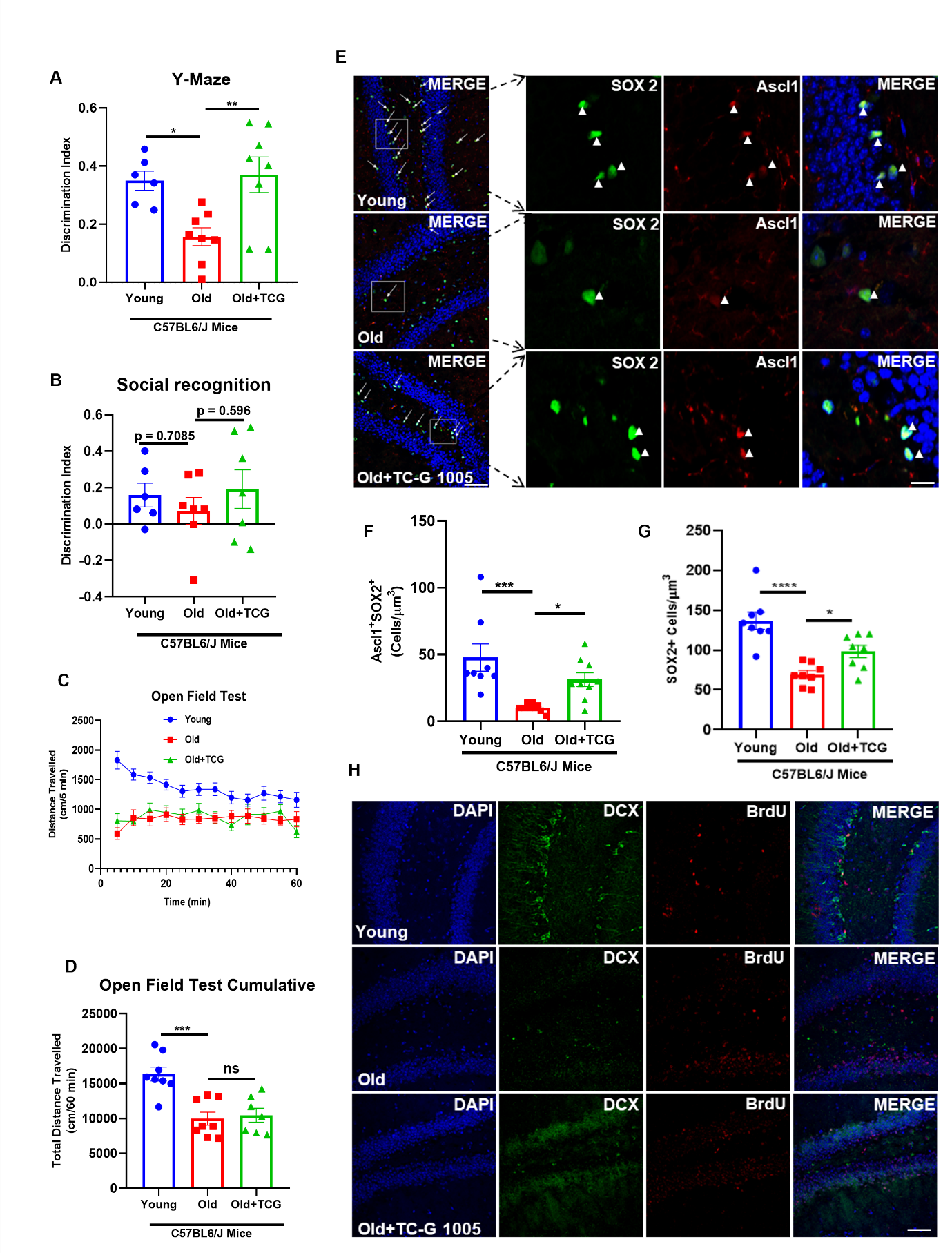

**Figure S8. (A)** Bar graph showing the discrimination index between a novel and familiar arm in the Y-maze test after 28 days of chronic subcutaneous injection of TC-G 1005. Data were plotted as mean ± SEM, Ordinary one-way ANOVA followed by Dunnett’s multiple comparisons test, (*p = 0.0186; Young vs Old, **p = 0.0055 Old vs Old + TCG 1005); n = 6-8/ group. **(B)** Representative bar graph showing an improvement trend in discrimination index between a novel and familiar mice in the social recognition test after 28 days of chronic subcutaneous injection of TC-G 1005. Data were plotted as mean ± SEM, Ordinary one-way ANOVA followed by Dunnett’s multiple comparisons test, (p = 0.7085; Young vs Old, p = 0.596 Old vs Old + TCG 1005); n = 6-8/ group. **(C & D)** Effect of TC-G 1005 on locomotor activity, total distance traveled were recorded for 60 min at 5-minute intervals and no significant change was observed in the locomotor activity. Data were plotted as mean ± SEM, Ordinary one-way ANOVA followed by Dunnett’s multiple comparisons test, (***p = 0.0003; Young vs Old, p = ns; Old vs Old + TCG 1005); n = 6-8/ group. **(E)** Representative images of dentate gyrus showing increase in Ascl 1 and increase in SOX2 population in old mice treated with TC-G 1005 suggesting activation of quiescent hiNSC and and therefore increase in stem cell pool. (Scale bar 49 µm). **(F)** Bar graph depicting quantitation of Ascl 1+SOX2+ cells as shown in (H). Data were plotted as mean ± SEM, Ordinary one-way ANOVA followed by Dunnett’s multiple comparisons test, (***p = 0.0007; Young vs Old, *p = 0.0421 Old vs Old + TCG 1005); n = 8-9/ group. **(G)** Bar graph depicting quantitation of SOX2+ cells as shown in (H). Data were plotted as mean ± SEM, Ordinary one-way ANOVA followed by Dunnett’s multiple comparisons test, (****p < 0.0001; Young vs Old, *p = 0.0371 Old vs Old + TCG 1005); n = 8-9/ group. **(H)** Representative images of dentate gyrus showing no effect on DCX expression in old mice treated with TC-G 1005 suggesting no effect on differentiation. (Scale bar 49 µm)

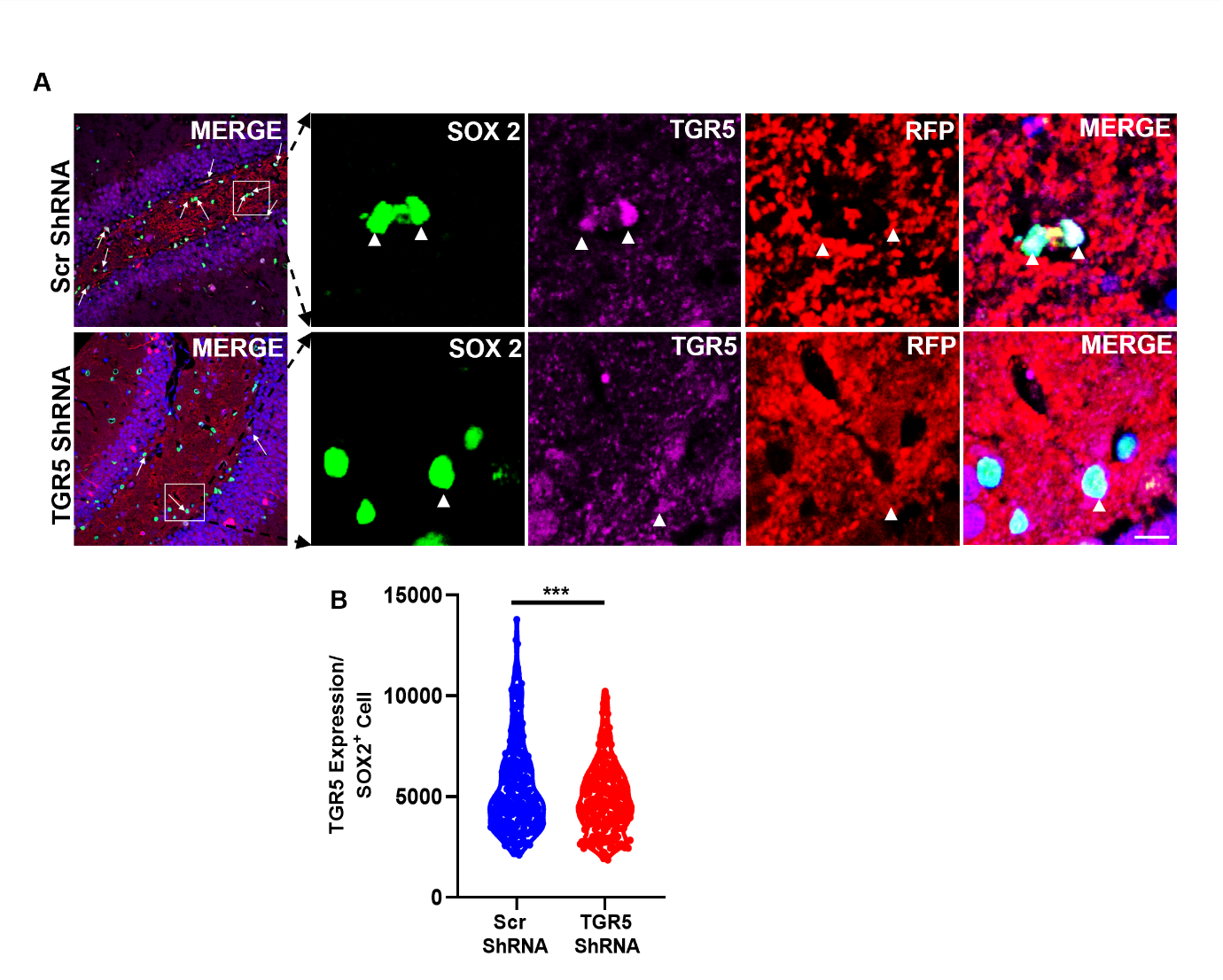

**Figure S9. (A)** Representative immunofluorescent images of scr- or AAV-TGR5-shRNA transduced Dentate gyrus depicting decreased TGR5expression of upon knockdown. (Scale bar 49 µm)**. (B)** Quantitation of TGR5 expression upon knockdown as shown in (A). Data were plotted as mean± SEM, unpaired t-test, t = 3.321, ***p = 0.0010, Scr-ShRNA vs TGR5-ShRNA n = 3 independent experiment

Antibodies used in Western Blotting.

| **Antibody** | **Source** | **Dilution** |
| --- | --- | --- |
| Anti-LC3β | Sigma, L7543 & Cell Signalling Technologies #12741 | 1:1000 |
| Anti p53 | Thermo fisher scientific MA5− 12571 | 1:1000 |
| Anti-Beclin-1 (D40C5) | Cell Signalling Technologies #3495 | 1:1000 |
| Anti-PCNA (PC10) | Cell Signalling Technologies #2586 | 1:1000 |
| Anti-p16 INK4A (E5F3Y) | Cell Signalling Technologies #29271 & Thermo fisher scientific PA5-20379 | 1:1000 |
| Anti-LC3A/B (D3U4C) | Cell Signalling Technologies #12741 | 1:1000 |
| Anti-ASCL1 (E5S4Q) | Cell Signalling Technologies #10585 | 1:1000 |
| Anti-FoxO3a (75D8) | Cell Signalling Technologies #2497 | 1:1000 |
| Anti-SQSTM1/p62 | Cell Signalling Technologies #5114 | 1:1000 |
| Anti Total-EGFR | Thermo fisher scientific PA1-1110 | 1:1000 |
| Anti-pEGFR (pTyr1068) | Cell Signalling Technologies #3777 | 1:1000 |
| Anti TGR5 | Abcam ab72608 | 1:1000 |

Antibodies used in Immunofluorescence.

| **Antibody** | **Source** | **Dilution** |
| --- | --- | --- |
| Anti-BrdU | Millipore MAB3510 | 1:500 |
| Anti-SOX2 | Thermo fisher scientific PA1-094 & 14-9811-82 | 1:500 |
| Anti-doublecortin | Abcam ab18723 | 1:500 |
| Anti-TGR5 | Abcam ab72608 | 1:500 |
| Anti-Ascl1 | Santa cruz sc-374104 | 1:200 |
| Anti-LAMP2 | Thermo fisher scientific PA1-655 | 1:500 |
| Anti-Nestin | Novus Biologicals  NB100-1604 | 1:500 |
| Anti-p16 | Santa Cruz sc-1661 | 1:500 |
| Anti-tGFP | Origene TA150041 & TA150075 | 1:500 |
| Anti-GFAP | Thermo fisher scientific  13-0300 | 1:500 |
| Anti-MAP2 | Sigma M9942 | 1:500 |
| Anti RFP | Thermo fisher scientific MA5-15257 | 1:500 |
